# Preclinical validation of a second generation leishmanization vaccine against vector transmitted fatal visceral leishmaniasis

**DOI:** 10.1101/2020.11.18.388553

**Authors:** Subir Karmakar, Nevien Ismail, Fabiano Oliveira, James Oristian, Wen Wei Zhang, Swarnendu Kaviraj, Kamaleshwar P. Singh, Abhishek Mondal, Sushmita Das, Krishna Pandey, Parna Bhattacharya, Greta Volpedo, Sreenivas Gannavaram, Monika Satoskar, Sanika Satoskar, Rajiv M. Sastry, Claudio Meneses, Shinjiro Hamano, Pradeep Das, Greg Matlashewski, Sanjay Singh, Shaden Kamhawi, Ranadhir Dey, Jesus G. Valenzuela, Abhay Satoskar, Hira L. Nakhasi

**Affiliations:** Division of Emerging and Transfusion Transmitted Diseases, CBER, FDA, Silver Spring, Maryland 20993, USA; Vector Molecular Biology Section, Laboratory of Malaria and Vector Research, National Institute of Allergy and Infectious Diseases, NIH, Rockville, Maryland 20852, USA; Department of Microbiology and Immunology, McGill University Montreal Canada, H3A 2B4; Gennova Biopharmaceuticals, Hinjawadi Phase II, Pune, Maharashtra 411057, India; Department of Parasitology, Institute of Tropical Medicine (NEKKEN), Nagasaki University, Nagasaki, Japan; Department of Pathology and Microbiology, Ohio State University, Columbus, Ohio, 43210, USA; Rajendra Memorial Research Institute of Medical Sciences, Patna, 800 097, India

## Abstract

Visceral Leishmaniasis (VL) is fatal if untreated. There is no licensed vaccine available against human leishmaniasis. We recently demonstrated protection in mice against *L. major* infection using a CRISPR genome edited attenuated *Leishmania major* strain (*LmCen*^*−/−*^). Here, as a pre-clinical step, we evaluated the protective efficacy of *LmCen*^*−/−*^ against VL induced by sand fly transmitted *Leishmania donovani* in hamsters. Intradermal immunization of hamsters with *LmCen*^*−/−*^ did not develop any lesion; while still priming a pro-inflammatory immune response. When challenged with *L. donovani* either by intradermal needle injection or by infected sand flies, *LmCen*^*−/−*^-immunized hamsters were protected, not showing spleen or liver pathology averting VL fatality compared to control animals. Spleen cells from *LmCen*^*−/−*^ immunized and infected sand fly challenged hamsters produced significantly higher Th1-associated cytokines and chemokines including IFN-γ and TNF-α, and significantly reduced expression of the anti-inflammatory cytokines IL-10 and IL-21, compared to non-immunized challenged animals. We further developed a GLP-grade *LmCen*^*−/−*^ which showed equal protection as laboratory-grade *LmCen*^*−/−*^ parasites in hamsters. Importantly, GLP-grade *LmCen*^*−/−*^ parasites also induced a proinflammatory immune response in the PBMCs isolated from healthy people living in non-endemic and endemic for VL as well as cured VL people living in endemic region. Together, this study demonstrates that the *LmCen*^*−/−*^ parasites are safe and efficacious against VL and it is a strong candidate vaccine to be tested in a human clinical trial.

## INTRODUCTION

Leishmaniasis is a complex disease caused by *Leishmania* infected sand fly bites. Among three major medical manifestation of leishmaniasis, visceral leishmaniasis (VL) is the most severe form of the disease and is fatal if untreated (***1***). Since curative drugs are toxic and often leads to drug resistance cases (***2***), a vaccine would be the best alternative. Patients who recover from leishmaniasis including VL develop protective immunity against re-infection, suggesting a road map for successful vaccine candidate where infection without pathology can induce a protective immunity (***3, 4***). However, currently there is no licensed human vaccine available for leishmaniasis.

The immunological protective mechanism in VL is complex. Cell mediated immunity is crucial for induction of host protective immune response, particularly Th1 immune response characterized by the production of IL-12 and IFN-γ (***5, 6***). In contrast, disease progression is associated with a dominant Th2 type immune response, mediated by IL-10 and IL-21 (***7***). Although, this Th1/Th2 dichotomy is clear in cutaneous leishmaniasis, it is not as defined in VL (***8, 9***). A study of in vivo cytokine profile in VL patients showed elevated levels of IL-10 and IFN-γ expression, while the levels of IL-10 decreased markedly with resolution of disease (***10, 11***). In addition, successful immunity against leishmaniasis also involves chemokines and chemokine receptors for the recruitment of immune effector cells to the infected sites, induction of genes coding for transcription factors, and finally activation of “classically activated” (M1) macrophages for the elimination of intracellular parasites through triggering an oxidative burst (***12, 13***).

Though several recombinant protein and DNA vaccines with or without adjuvant showed promising results in animal model, they failed to achieve satisfactory results in clinical trials (***11, 14, 15***). Among the several vaccination strategies attempted against leishmaniasis, infection with low dose of live wild type *Leishmania* promastigotes (leishmanization), was the only successful immunization strategy for CL in humans (***16–18***). However, under the current regulatory environment such practice is not acceptable due to safety concerns (***19, 20***). Therefore, live attenuated *Leishmania* parasites that are nonpathogenic and provide a complete array of antigens of a wild type parasite should induce the same protective immunity as “leishmanization” and thus would be an effective vaccine candidate (***21***).

Prior studies from our group demonstrated that live attenuated *centrin* gene deleted *Leishmania donovani* parasite (*LdCen*^*−/−*^) protects against *L. donovani* challenge in various pre-clinical animal models of VL (***22–24***). Centrin is a calcium binding protein and essential in the duplication of centrosomes in eukaryotes including *Leishmania* (***25, 26***). It is important to note that *centrin* gene deficient *Leishmania* parasites are attenuated only in intracellular amastigote stage and can be grown in promastigote culture (***25***).

Several studies have documented that exposure to wild type *L. major* parasites which causes localized self-resolving infection confers cross-protection against VL in various animal models (***27, 28***). Epidemiological evidence suggested that cutaneous infection with *L. major* cross-protects against visceral leishmaniasis in humans (***29***). All these findings suggest that live attenuated dermotropic *Leishmania* parasites could be a promising vaccine against CL as well as VL and would be safer than using a viscerotropic live attenuated *L. donovani* strain. We have recently reported that *centrin* gene deleted *L. major* (*LmCen*^*−/−*^) parasites are safe and protect against cutaneous infection with wild type *L. major* in a mice model (***30***). This represented a major breakthrough because using CRISPR, it was possible to engineer *LmCen*^*−/−*^ without any antibiotic resistance marker genes making it compliant for human vaccine trials.

In the present study, we tested the safety, immunogenicity and efficacy of the live attenuated dermotropic *LmCen*^*−/−*^ parasites as a vaccine candidate against *L. donovani* infection initiated by natural vector bites using preclinical hamster model considered to be a gold standard for VL disease. In addition, we tested immunogenicity of GLP-grade *LmCen*^*−/−*^ parasites in the PBMCs isolated from healthy people living in VL endemic and non-endemic regions and Cured VL individuals from endemic regions.

## RESULTS

### Immunization with live attenuated *LmCen*^*−/−*^ parasites does not cause skin pathology in hamsters

To evaluate the safety of *LmCen*^*−/−*^ parasites as a vaccine, hamsters were injected intradermally with either *LmCen*^*−/−*^ promastigotes or wild type *L. major* (*LmWT*) and monitored for lesion development and parasite burden (**Fig. 1A**). *LmCen*^*−/−*^ injected hamsters did not develop any visible lesions up to 49-days of post-injection (**Fig. 1B, C**). In contrast, hamsters injected with *LmWT* parasites, develop ear lesions within 15 days of parasite injection that progressively increased in size (**Fig. 1B, C**). At three days post-injection, the parasite load in the ears and dLNs was similar in both *LmWT* and *LmCen*^*−/−*^ injected hamsters (**Fig. 1D, E**). At 15 days post-inoculation, we started to observe a significant difference in parasite load between these groups with *LmWT* injected-hamsters displaying a higher parasite load as compared to lower numbers in *LmCen*^*−/−*^-injected hamsters (**Fig. 1D, E**). The difference in parasite load between these two groups was greater at days 28 and 49 (**Fig. 1D, E**). Importantly, two of six *LmCen*^*−/−*^ injected hamsters cleared the parasites by 28 days post injection from ear. Furthermore, at day 49, no viable parasites were recovered from the ears or dLNs of hamsters injected with *LmCen*^*−/−*^ parasites (**Fig. 1D, E**).

**Fig. 1.**
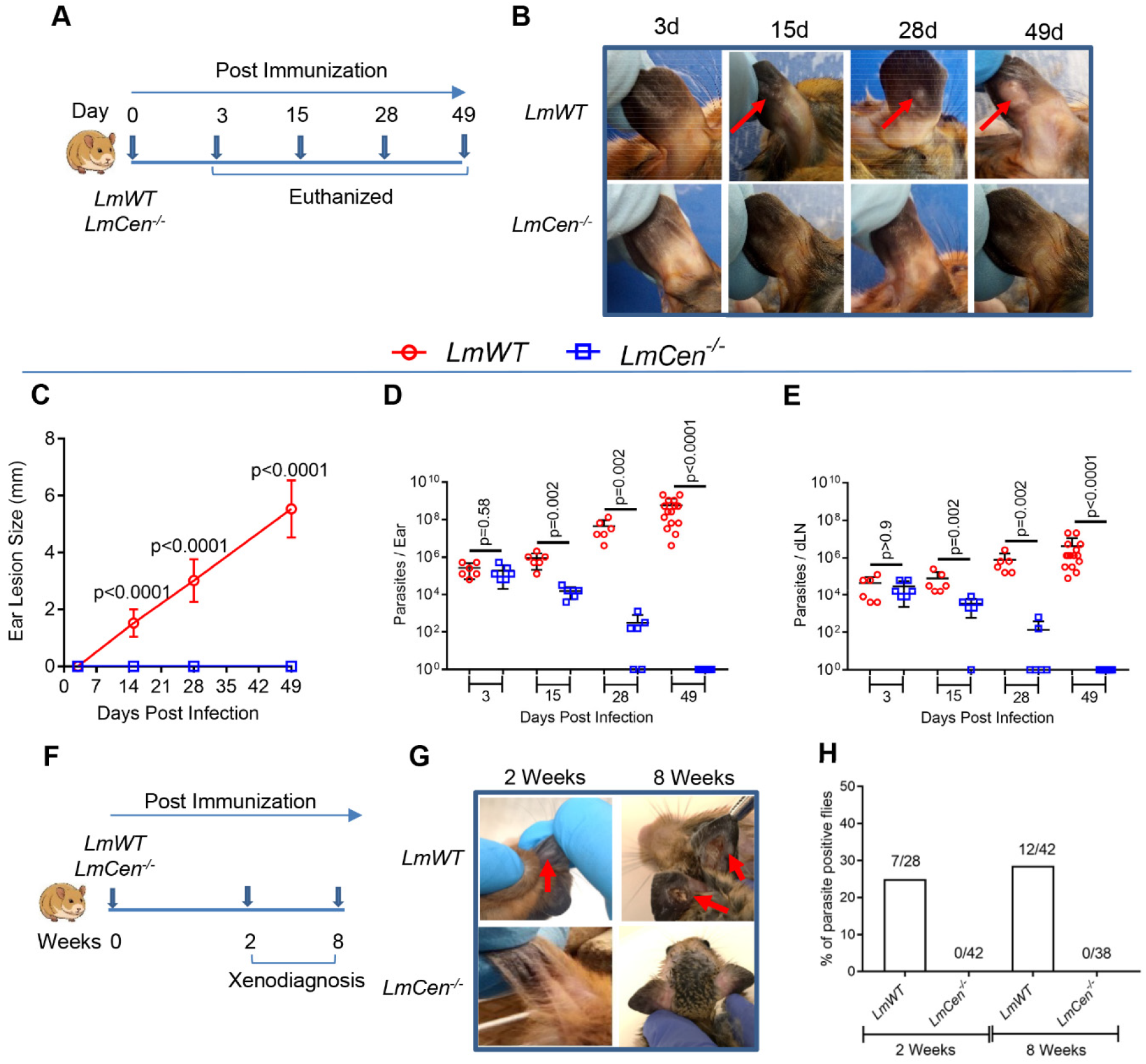
Live attenuated *LmCen*^*−/−*^ parasites do not cause any pathology in hamster model. (**A**) Schematic representation of experimental plan. Lesion size was monitored every week in hamsters injected with 10^6^-total stationary phase either *LmWT* or *LmCen*^*−/−*^ parasites by intradermal (ID) injection. (**B**) Photographs of representative ears of *LmWT* and *LmCen*^*−/−*^ immunized hamsters at indicated days post inoculation. Red arrows indicate the lesion development. (**C**) Ear lesion diameters were measured at indicated days post inoculation. Results (SD) are representative cumulative effect of two (3d, 15d and 28d) to three (49d) independent experiments, 1 ear, total 6-33 hamsters per group (p values were determined by Mann Whitney two-tailed test). (**D** and **E**) Parasites load in the ear (D) and draining lymph node (dLN) (E) of *LmWT* and *LmCen*^*−/−*^ immunized hamsters (n=6 for 3, 15, 28 day and n=15 for 49 day for both the group of hamsters) were determined by serial dilution assay at indicated time points. Results (Mean ± SD) represent cumulative effect of two (3d, 15d and 28d) to three (49d) independent experiments (p values were determined by Mann Whitney two-tailed test). (**F**) Schematic representation of xenodiagnoses to determine infectiousness of immunized hamsters for sand flies. (**G**) Photographs of representative ears of *LmWT* and *LmCen*^*−/−*^ infected hamsters for xenodiagnoses at 2- & 8-weeks post inoculation. Red arrows indicate the lesion development. (**H**) After exposed with infected hamsters (n=6/group), blood fed flies were isolated, and parasite positive flies were identified after dissection of flies isolated from both *LmWT* and *LmCen*^*−/−*^ infected groups. Results (Mean ± SD) are representative one experiment.

To rule out the possibility that hamsters injected with *LmCen*^*−/−*^ parasites can serve as reservoirs of these parasites to sand flies, we performed a xenodiagnosis test using non-infected sand flies at 2- and 8-weeks post injection (**Fig. 1F**), timepoints where parasites were most numerous or undetectable, respectively, after injection with *LmCen*^*−/−*^ parasites (**Fig. 1G**). *L. longipalpis* sand flies fed on hamsters injected with *LmCen*^*−/−*^ parasites were all negative at both tested timepoints (**Fig. 1H**). In contrast, 25 to 30% sand flies exposed to *LmWT* injected animals were *Leishmania*-positive (**Fig. 1H**). These results demonstrate that live *LmCen*^*−/−*^ parasites do not establish an infection in vector sand flies.

### Immunization of *LmCen*^*−/−*^ parasites does not cause lesions in immune suppressed animals

To assess the safety of *LmCen*^*−/−*^ parasites, we investigated lesion development and survival of *LmCen*^*−/−*^ parasites in immune-suppressed animals treated with dexamethasone (DXM) (**Fig. 2A**). Immune-suppressed hamsters injected with *LmCen*^*−/−*^ parasites showed no lesions at the inoculation site (ear) compared to the ulcerative lesions that developed in *LmWT* injected hamsters four weeks after immune-suppression (**Fig. 2B, C**). Only three of the 12 *LmCen*^*−/−*^ injected/immune-suppressed hamsters had parasites in the inoculated ear (**Fig. 2D**) and in the dLN (**Fig. 2E**). As expected, all the *LmWT*-injected hamsters had significantly higher parasite loads in the ear (**Fig. 2D**) and in the dLN (**Fig. 2E**) compared to *LmCen*^*−/−*^-injected animals (±immune-suppressed). To investigate whether *LmCen*^*−/−*^ parasites isolated from immunosuppressed animals reverted to the wild type genotype, we performed PCR analysis of their genomic DNA. We confirmed the absence of the *centrin* gene in the parasites isolated from all the three *LmCen*^*−/−*^-injected immune-suppressed hamsters (**Fig. 2F**, lane 2, 3, 4 and 5, red arrow). We further examined whether the *LmCen*^*−/−*^ parasites recovered from immune-suppressed hamsters had regained virulence by testing them in human monocyte-derived macrophages (hMDM). After 144h infection, *LmCen*^*−/−*^ parasites were mostly cleared from the hMDM whereas the number of *LmWT* parasites reached >6 parasites/hMDM (**Fig. 2G, H**). Collectively, these results demonstrate that the attenuated *LmCen*^*−/−*^ parasites are safe, unable to revert to the wild type form even in an immune-suppressed condition.

**Fig. 2.**
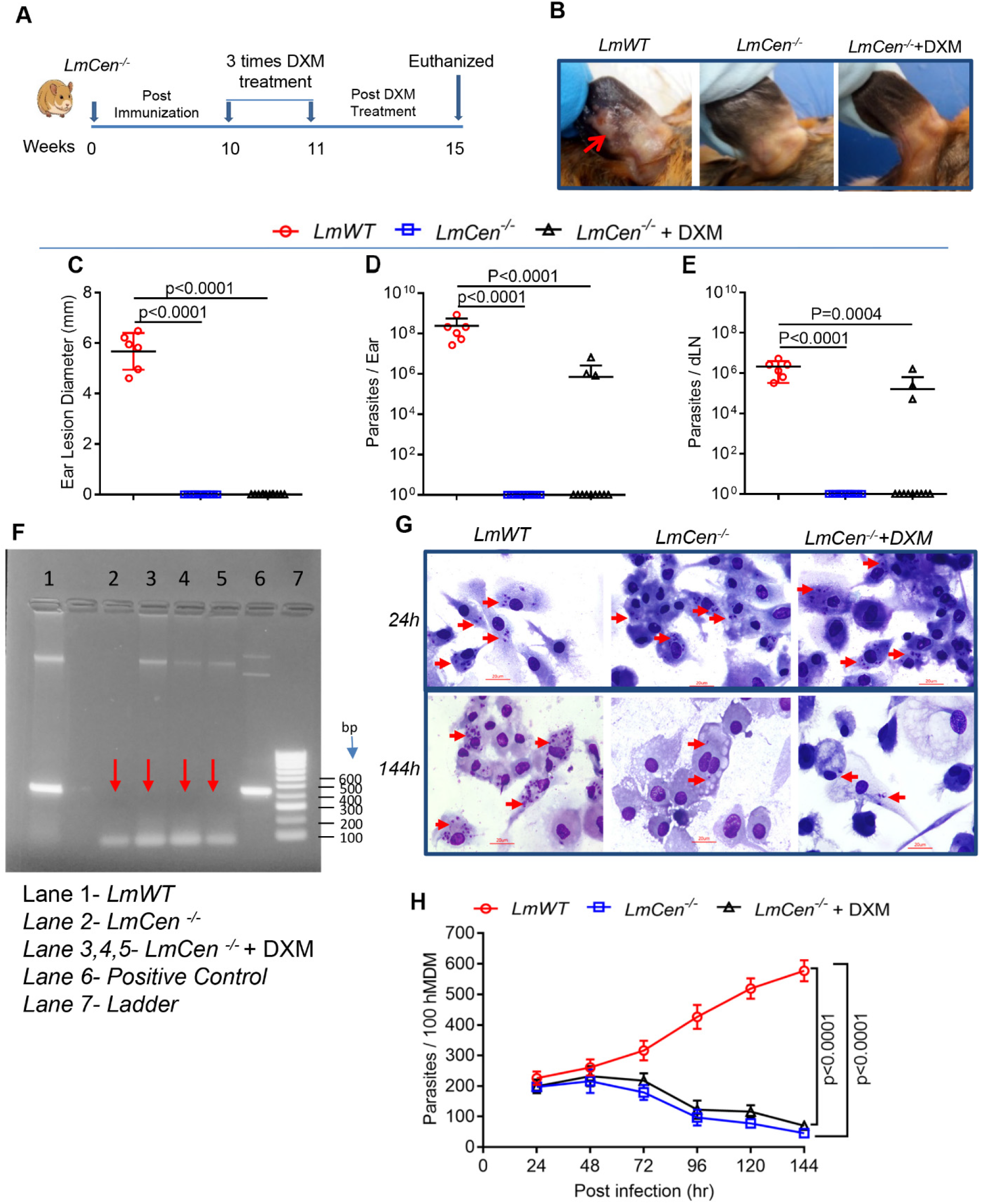
*LmCen*^*−/−*^ does not cause any lesions even in immune suppressed animals. (**A**) Schematic representation of immune-suppression by DXM treatment of *LmCen*^*−/−*^ immunized hamsters. (**B**) Photographs of representative ears of *LmWT*, *LmCen*^*−/−*^ and *LmCen*^*−/−*^+DXM treated hamsters. Red arrow indicates the lesion development. (**C**) Ear lesion diameters were measured after 4 weeks of DXM treatment (total 15 weeks post parasite infection) in *LmWT* (n=6) and *LmCen*^*−/−*^ (n=12) and *LmCen*^*−/−*^+DXM (n=12) treated hamsters. Results (Mean ± SD) represent cumulative effect of two independent experiments, 1 ear, total 6-12 hamsters per group (p values were determined by Mann Whitney two-tailed test). (**D** and **E**) Parasite load in the inoculated ear (D) and dLN (E) of each group of hamsters (*LmWT,* n=6; *LmCen*^*−/−*^, n=12; and *LmCen*^*−/−*^ + DXM, n=12) were determined by limiting dilution assay. Results (Mean ± SD) represent cumulative effect of two independent experiments (p values were determined by Mann Whitney two-tailed test). (**F**) 1% Agarose gel electrophoresis results for the characterization of *LmCen*^*−/−*^ parasites isolated from *LmCen*^*−/−*^ plus DXM treated group using *L. major centrin* gene specific primers. Lane-1, PCR results from the genomic DNA of parasites isolated from *LmWT*, Lane-2, PCR results from the genomic DNA of parasites isolated from *LmCen*^*−/−*^ treated group, Lane-3,4,5 PCR results from the genomic DNA of parasites isolated from *LmCen*^*−/−*^ + DXM treated group, Lane-6 PCR results from the plasmid DNA containing *centrin* gene as a positive control. Lane 4, 100 bp DNA ladder (Bioline). Red arrow indicates the absence of main product bands (*centrin* gene) of 450bp in Lane-2,3,4 and 5. (**G** and **H**) Photographs (G) of human monocyte derived macrophages (hMDM) at 24h and 144h post-infection with parasites isolated from *LmCen*^*−/−*^ plus DXM treated group as well as *LmWT* and *LmCen*^*−/−*^ parasites respectively. The number of amastigotes (red arrow) was determined microscopically up to 144h post infection. The data (Mean ± SD) are represented as the number of parasites per 100 hMDM (H) of one independent experiment (p values were determined by Unpaired two-tailed t test).

### Immunization with *LmCen*^*−/−*^ induces a local and systemic pro-inflammatory immune response

We examined the local and systemic immune response at the inoculation site (ear) and spleen, respectively, of *LmCen*^*−/−*^-immunized hamsters. We observed a higher expression of transcripts related to pro-inflammatory or Th1 type cytokines (IFN-γ, TNF-α, IL-1β and IL-6) in the ear of *LmCen*^*−/−*^-immunized animals compared to animals injected with *LmWT* (**Fig. 3A, B**; Fig S1A). Furthermore, the expression of anti-inflammatory (IL-4 and IL-21) and regulatory (IL-10) cytokine transcripts were significantly lower in *LmCen*^*−/−*^-immunized animals compared to the *LmWT*-injected group (**Fig. 3A, B**). Additionally, the transcription factor T-bet transcript, which is involved in the regulation of a pro-inflammatory response, was significantly higher in the *LmCen*^*−/−*^ immunized group (**Fig. 3A;** Fig S1A). Conversely, transcription factors GATA3 and Foxp3 transcripts, involved in Th2 or anti-inflammatory type cytokine responses, respectively, were higher in *LmWT* injected group (**Fig. 3A**; Fig S1A). Regarding pro-inflammatory chemokines, the expression of the CCL5 and CXCL9 transcripts were significantly higher in the *LmCen*^*−/−*^ immunized hamsters compared to *LmWT* injected animals (**Fig. 3A**; Fig. S1A). *LmWT* injected animals had a higher expression CCL4 transcript compared to *LmCen*^*−/−*^ immunized animals (**Fig. 3A;** Fig. S1A). The expression profile of transcripts of markers of alternatively activated M2 macrophages were significantly lower (Arg-1 and CCL17) in the *LmCen*^*−/−*^ immunized group compared to *LmWT* injected hamsters (**Fig. 3A**; Fig. S1A). Taken together, these results suggest that *LmCen*^*−/−*^ induce a pro-inflammatory/ Th1 dominant immune response, while *LmWT* induce anti-inflammatory disease promoting response.

**Fig. 3.**
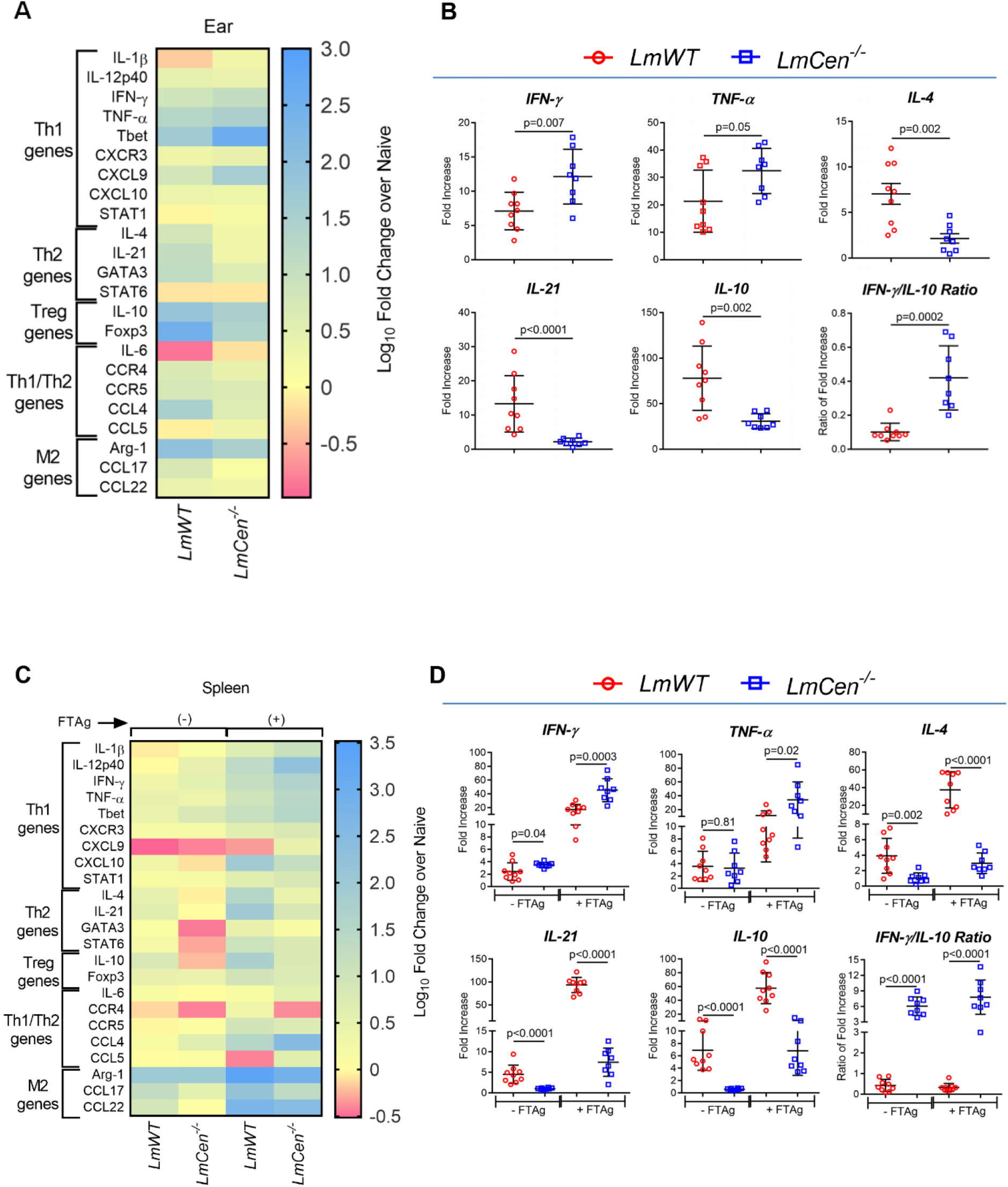
*LmCen*^*−/−*^ immunization induces pro-inflammatory immune response. (**A** and **C**) Heat map showing the differential gene expression in the ear (A) (without antigen re-stimulation) and spleen (C) (with or without 24h of *L. major* freeze thaw antigen re-stimulation;±FTAg) of *LmWT* infected and *LmCen*^*−/−*^ immunized hamsters at 7 weeks post inoculation. Down-regulation & up-regulation of the transcripts are shown in blue, yellow and pink respectively. Transcripts are annotated in the left side according to their general functions. For heat map, results were shown as log10 fold change over naive hamster. (**B** and **D**) Expression profile of IFN-γ, TNF-α, IL-4, IL-21, IL-10 and IFN-γ/IL-10 Ratio in the ear (B) and spleen (D) which was evaluated by RT-PCR. The data were normalized to γ-Actin expression and shown as the fold change relative to age matched naive hamster. Results (Mean ± SD) represent cumulative effect of two independent experiments (n=9 for *LmWT and n=8 for LmCen*^*−/−*^) (p values were determined by Mann-Whitney two-tailed test).

For systemic immune responses in the spleen after splenocyte stimulation with freeze-thaw *Leishmania* antigen (FTAg), the expression profile of transcripts for Th1 type cytokines (IFN-γ, TNF-α, IL-1β, IL-12p40 and IL-6) transcription factors (T-bet and STAT1) and chemokines/their ligands (CXCL9) were significantly higher in the *LmCen*^*−/−*^ immunized animals compared to *LmWT* injected hamsters (**Fig. 3C, D**; Fig. S1B). However, splenocytes from *LmWT* infected hamsters had a significantly higher expression of both, anti-inflammatory (IL-4, IL-21, GATA3 and STAT6) as well as regulatory (IL-10 and Foxp3) transcripts compared to *LmCen*^*−/−*^ immunized group (**Fig. 3C, D**; Fig. S1B). The ratio of IFN-γ to IL-10 was significantly higher in both the ear and spleen of the *LmCen*^*−/−*^ immunized group compared to *LmWT* injected animals (**Fig. 3B** and **3D**). These results collectively suggest that *LmCen*^*−/−*^ immunization induces a pro-inflammatory environment in the ear and spleen of immunized animals.

### Immunization with *LmCen*^*−/−*^ parasites protects hamsters against needle challenge with *L. donovani*

Previous studies have suggested that cutaneous infection with *L. major* cross-protects against visceral leishmaniasis in humans (*29*). We, therefore, investigated the efficacy of immunization with *LmCen*^*−/−*^ parasites against visceral leishmaniasis induced by intradermal injection of *L. donovani* parasites in hamsters (**Fig. 4A**). There was a significant protection in *LmCen*^*−/−*^ immunized hamsters as compared to non-immunized-infected animals as demonstrated by the smaller size of the spleen in *LmCen*^*−/−*^ immunized hamsters compared to the enlarged spleen in non-immunized animals, 12-month post-challenge (**Fig. 4B & C**). Numerous *L. donovani* amastigotes were observed in smears from spleen tissue of non-immunized group compared to *LmCen*^*−/−*^ immunized hamsters (**Fig. 4C**). Furthermore, non-immunized hamsters showed progressive weight loss beginning at nine months post-challenge (Fig. S2A). Additionally, the spleen weight in the non-immunized-challenged hamsters was significantly higher compared to *LmCen*^*−/−*^ immunized-challenged animals, three months post challenge (**Fig. 4B**). Importantly, all the non-immunized hamsters exhibited severe splenomegaly nine months post challenge. Since successful resistance to *L. donovani* in hepatic tissues is reflected by mature granuloma formation (***31***), we also examined liver sections from immunized and non-immunized hamsters 9 months post-challenge. Histopathological analysis of liver from non-immunized animal revealed heavily parasitized Kupffer cells (black arrows) with few infiltrating lymphocytes resembling an immaturely formed granuloma (Fig. S2B). In contrast, *LmCen*^*−/−*^ immunized hamsters exhibited well-formed granulomas comprised of concentric mononuclear cells (Fig. S2B).

**Fig. 4.**
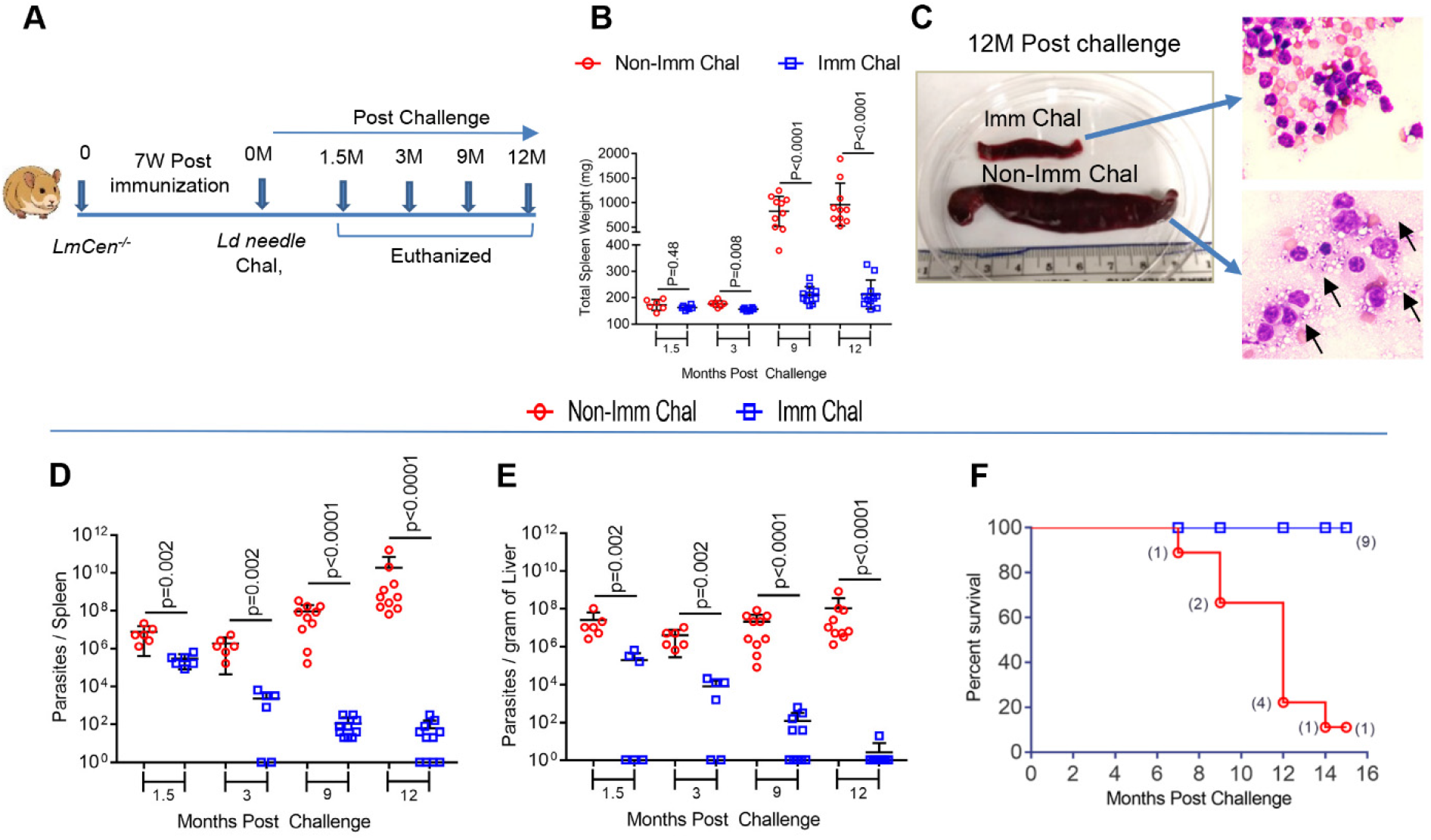
Immunization with *LmCen*^*−/−*^ parasites protects hamsters against needle challenge with *L. donovani*. (**A**) Schematic representation of experimental plan to determine the cross-protective efficacy of *LmCen*^*−/−*^ parasites against *L. donovani* via needle challenge. (**B** and **C**) Spleen weight (B) of *LmCen*^*−/−*^ immunized (Imm Chal) (n=6 for 1.5 and 3 months and n=10 for 9 and n=11 for 12 months) and age matched non-immunized (Non-Imm Chal) hamsters (n=6 for 1.5 and 3 months and n=10 for 9 and 12 months) after various periods of post needle challenge (1.5, 3, 9 & 12 month) with *L. donovani.* Results (Mean ± SD) are representative of cumulative effect of two independent experiments (p values were determined by Mann Whitney two-tailed test). A representative spleen sample (C, left panel) of *LmCen*^*−/−*^ immunized (Imm Chal) and non-immunized (Non-Imm Chal) hamsters after 12 months post needle challenge. Spleen size of the both groups of hamsters was shown in centimeter and H&E stained stamp smears (right panel) from respective spleen were shown where black arrow indicates intracellular *L. donovani* amastigotes. (**D** and **E**) Parasite load were determined by limiting dilution after various periods post challenge and expressed as number of parasites per Spleen (D) and per gram of Liver (E). Results (Mean ± SD) represent cumulative effect of two independent experiments (p values were determined by Mann Whitney two-tailed test). (**F**) Kaplan-Meier survival curves comparing survivability of *LmCen*^*−/−*^ immunized (n=9) hamsters with age matched non-immunized *L. donovani* challenged (n=9) hamsters. Results are representative of one experiment.

Analysis of the parasite load revealed significant control of parasite numbers in the spleen (**Fig. 4D**) and liver (**Fig. 4E**) of *LmCen*^*−/−*^ immunized hamsters compared to non-immunized-infected animals at all timepoints tested. Immunized hamsters showed ~1.5 log-fold reduced parasite burden in spleen and liver as early as 1.5-month post challenge. By 12 months post-challenge, the parasite burden was reduced by ~5 log-fold for the spleen (**Fig. 4D**) and ~12 log-fold for the liver (**Fig. 4E**). Of note, at 12 months post challenge, 36% (4 of 11) of the spleens and 90% (10 of 11) of the livers from immunized animals had undetectable numbers of viable parasites. All the *LmCen*^*−/−*^ immunized hamsters survived the lethal challenge of *L. donovani* and remained healthy up to 14-months post-challenge, the end of our study period, whereas eight out of nine non-immunized challenged hamsters died within 14-months (**Fig. 4F**). Taken together, these data demonstrate that *LmCen*^*−/−*^ elicits protection against needle infection in a hamster model of VL.

### *LmCen*^*−/−*^-immunized hamsters induce a pro-inflammatory/Th1 type of immune response upon challenge with wild type *L. donovani*

To characterize the immune correlates of protection, we measured the gene expression profile *ex-vivo* for the challenged ear and after antigen re-stimulation for the spleen by qPCR following 1.5 months after needle challenge with virulent *L. donovani* parasites. Of the tested genes, the ear of immunized animals showed a significant increase in the expression profile of transcripts for pro-inflammatory cytokines (IFN-γ, IL-1β and IL-12p40), with a concomitant decrease in transcript expression of anti-inflammatory (IL-21) cytokines compared with non-immunized-challenged animals (**Fig. 5A, B**; Fig. S3A). Additionally, the expression of transcription factors, chemokines and chemokine receptors transcripts in the *LmCen*^*−/−*^ immunized challenged group were significantly higher for T-bet and CXCL9, related to Th1, and lower for GATA3, Foxp3, and STAT6 related to a Th2 response, and for CCR4 and CCR5 compared to non*-*immunized challenged animals at this time point (**Fig. 5A**; Fig. S3A). *LmCen*^*−/−*^ immunized challenged hamsters also had a significantly lower expression of transcripts (Arg-1, CCL17 and CCL22), characteristic of M2 macrophages, compared to non-immunized-challenged hamsters (**Fig. 5A**; Fig. S3A). Evaluation of the immune response in the spleen also showed a markedly increased expression of pro-inflammatory cytokine, chemokine and transcriptional factor transcripts (IFN-γ, TNF-α, IL-12p40, T-bet, STAT1 and CXCR3) with a significant decrease in anti-inflammatory genes (IL-21 and STAT6) in the immunized-challenged group compared to the non-immunized challenged group (**Fig. 5C, D**; Fig. S3B). Of Note, expression of M2 macrophage phenotype genes (CCL17) and chemokine and chemokine receptor transcripts (CCR4 and CCL4) were higher in the non-immunized challenged group compared to non-immunized challenged group (**Fig. 5C**; Fig. S3B). Importantly, the ratio of IFN-γ to IL-10 was significantly higher in both the ear and spleen of the immunized-challenged group compared to non-immunized challenged animals (**Fig. 5B** and **5D**). Collectively, these results indicate generation of a proinflammatory type of immune response following needle challenge in the *LmCen*^*−/−*^ immunized animals that confers protection against virulent *L. donovani*.

**Fig. 5.**
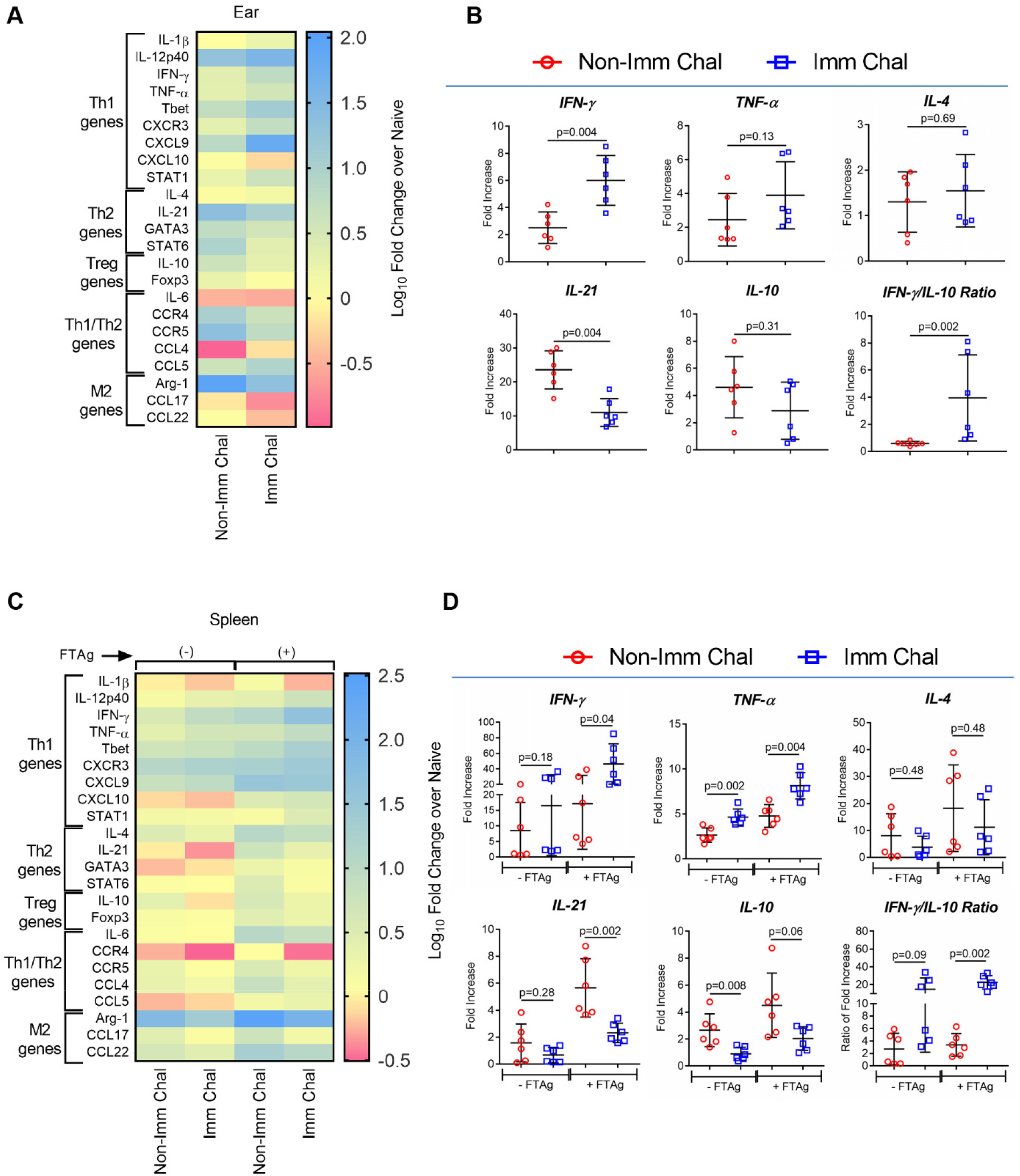
*LmCen*^*−/−*^ immunized hamster induces pro-inflammatory/Th1 type of immune response upon challenge with wild type *L. donovani*. (**A** and **C**) Heat map (left panel) showing the differential gene expression in the ear (A) (without antigen re-stimulation) and spleen (B) (with or without 24h of *L. donovani* freeze thaw antigen re-stimulation; ±FTAg) of age matched non-immunized (Non-Imm Chal) and *LmCen*^*−/−*^ immunized (Imm Chal) hamsters after 1.5 months of *L. donovani* challenge by needle injection (n=6/group). According to the general function, transcripts are annotated in the left side. Down-regulation & up-regulation of the transcripts are shown in pink, yellow and blue, respectively. For heat map, results were shown as log10 fold change over naive hamster. (**B** and **D**) Expression profile of IFN-γ, TNF-α, IL-4, IL-21, IL-10 and IFN-γ/IL-10 Ratio in the ear (B) and spleen (D) which was evaluated by RT-PCR. The data were normalized to γ-Actin expression and shown as the fold change relative to age matched naive hamster. Results (Mean ± SD) represent cumulative effect of two independent experiments (p values were determined by Mann-Whitney two-tailed test).

### *LmCen*^*−/−*^ immunization confers protection against sand fly-transmitted *L. donovani*

We determined the protective efficacy of *LmCen*^*−/−*^ immunization against infection initiated by sand fly transmitted *L. donovani*, considered a gold standard challenge. Sand flies were infected with *L. donovani* and the parasite load and quality of the infection in the insect gut was assessed before transmission (Fig. S4A). At 13 days after infection in the sand flies, the geometric mean parasite load (GMPL) per sand fly midgut was > 1.0 × 10^5^ *L. donovani* parasites, and the mean percent of metacyclic parasites per midgut was >80%, indicating a good quality infection in the sand fly (Fig. S4A). Seven weeks post immunization with *LmCen*^*−/−*^ parasites, hamsters were exposed to the bites of thirty *L. donovani*-infected sand flies in the contralateral ear (**Fig. 6A**). The mean number of fed flies per group of immunized and non-immunized hamsters were comparable (Fig. S4B). Immunization of hamsters with *LmCen*^*−/−*^ protected animals against visceral leishmaniasis. There was a significant reduction of parasite burden in both the spleen and liver of immunized hamsters compared to the progression of the infection in the non-immunize group (**Fig. 6B, C**). Parasite loads in immunized hamsters showed ~6.5 log-fold reduction in the splenic parasite burden and ~3.5 log-fold reduction in the liver at 9-month post challenge as compared to non-immunized hamsters (**Fig. 6B, C**). A ~10-log-fold reduction in the parasite load in the spleen (**Fig. 6B**), that was undetectable in the liver (**Fig. 6C**), was observed in immunized hamsters at 12-month post-challenge. A progressive infection was associated to a significant weight loss in all non-immunized hamsters compared to *LmCen*^*−/−*^ immunized hamsters beginning at 9-month post-challenge (**Fig. 6D**, **Fig. S4C**). Further, all the non-immunized-challenged hamsters exhibited severe splenomegaly compared to *LmCen*^*−/−*^ immunized or naïve animals at 9-12 months post challenge (**Fig. 6E**, **S4D**). Furthermore, numerous *L. donovani* amastigotes were observed in smears from spleen tissue of non-immunized in comparison to *LmCen*^*−/−*^ immunized hamsters (Fig. S4E). Most relevant, all the *LmCen*^*−/−*^ immunized hamsters survived the infected sand fly challenge and remained healthy up to 14 months post-challenge, the end of our study period, whereas all age matched non-immunized-challenged hamsters succumbed to virulent *L. donovani* challenge within 6-14 month (**Fig. 6F**).

**Fig. 6.**
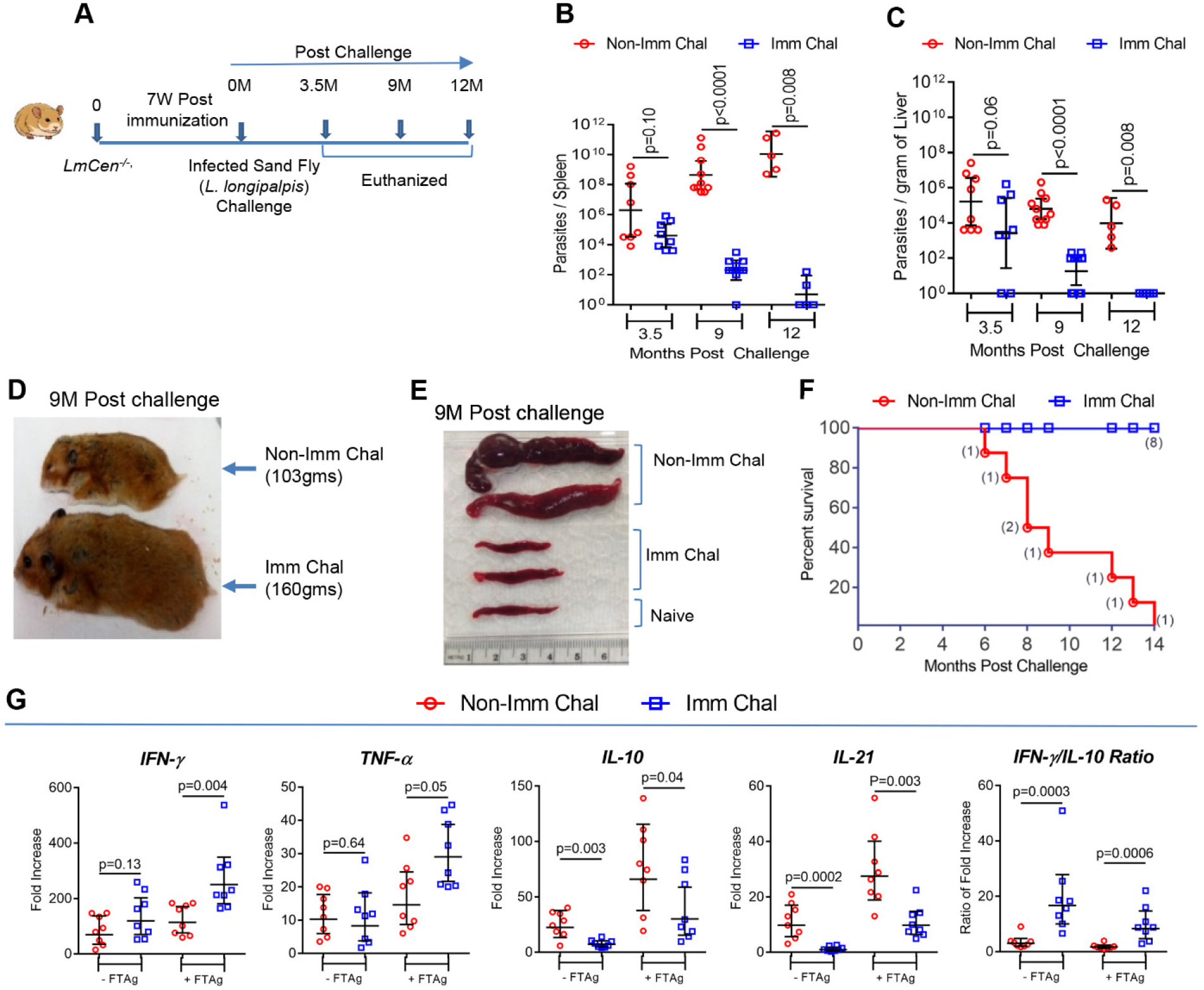
*LmCen*^*−/−*^ immunization confers protection against sand fly bite transmitted *L. donovani* infection in hamsters. (**A**) Schematic representation of experimental plan to determine the cross-protective efficacy of *LmCen*^*−/−*^ parasites against *L. donovani* infected sand fly (*Lutzomyia longipalpis*) challenge. (**B** and **C**) Parasite load in spleen (B) and liver (C) (n=8, n=10 and n=5 for 3.5M, 9M and 12M post challenge respectively of non-immunized and immunized challenged groups) were determined by limiting dilution at indicated time points following challenge and expressed as number of parasites per Spleen and per gram of Liver. Results represent (the geometric means with 95% Cl) cumulative effect of two independent experiments for 3.5- and 9-months post-challenge and one experiment for 12 months post-challenge (p values were determined by Mann-Whitney two-tailed test). (**D**) Picture showing the clinicopathologic features of VL in a representative sick age matched non-immunized hamster, compared with *LmCen*^*−/−*^-immunized healthy hamster after 9 months of sand fly-transmitted *L. donovani* challenge. (**E)** Photographs of representative two spleen samples of both *LmCen*^*−/−*^-immunized and non-immunized hamsters following 9-months post challenge as well as one age matched naive hamster is shown. Spleen size of all the hamsters was measured in centimeter. (**F**) Kaplan-Meier survival curves of *LmCen*^*−/−*^-immunized hamsters (Imm Chal; blue lines, n=8) following challenge with *L. donovani* infected sand flies and compared with age matched non-immunized challenged group (; red lines, n=8). (**H**) Immune response in the spleen ((with or with-out 24h *L. donovani* freeze-thaw antigen re-stimulation, ±FTAg) of *LmCen*^*−/−*^ immunized (n=8) and age matched non-Immunized (n=8) hamsters was determined following 3.5 months post *L. donovani* infected sand fly challenge. The expression of IFN-γ, TNF-α, IL-10 and IL-21was evaluated by RT-PCR. The ratio of IFN-γ/IL-10 expression in the spleen was also determined. Results were normalized to γ-actin expression and shown as the fold change relative to age matched naive hamster. Results (the geometric means with 95% Cl) represent cumulative effect of two independent experiments (p values were determined by Mann-Whitney two-tailed test).

To evaluate the protective immune response in *LmCen*^*−/−*^ immunized hamsters following vector transmission, cytokines expression was assessed in the spleen by RT-PCR at 3.5-month post challenge. Following antigen re-stimulation, the expression of the pro-inflammatory cytokine transcripts IFN-γ and TNF-α was significantly higher in *LmCen*^*−/−*^ immunized compared to non-immunized challenged hamsters (**Fig. 6G**). Concomitantly, there was a significantly higher expression of the anti-inflammatory cytokine transcripts IL-10 and IL-21 in the non-immunized-challenged group as compared to *LmCen*^*−/−*^ immunized animals (**Fig. 6G**). The ratio of IFN-γ to IL-10 transcript expression was significantly higher in *LmCen*^*−/−*^ immunized challenged animals compared to the non-immunized challenged group (**Fig. 6G**). Taken together, these data demonstrate that *LmCen*^*−/−*^ immunization mediated significant protection against VL initiated via the natural mode of parasite transmission with infected sand flies by inducing a protective pro-inflammatory immune response.

### GLP-grade *LmCen*^*−/−*^ parasites induce host protection in hamsters against challenge with wild type *L. donovani*

Advancing *LmCen*^*−/−*^ parasites as a vaccine for potential human clinical trials will require parasites grown under current Good Manufacturing Practices (cGMP). Here, we produced *LmCen*^*−/−*^ parasites in a bioreactor at small industrial scale under Good Laboratory Practices (GLP). GLP production of *LmCen*^*−/−*^ parasites is scalable and it can be transferred for cGMP production of *LmCen*^*−/−*^ parasites for future studies. Briefly, parasites were grown in a bioreactor 1L culture medium, and vials (1 million cells/mL, in 1.8mL volume) were prepared after 84-hours of culture when they reached a maximum cell density of 45-50 million cells per milliliter and stored in liquid nitrogen (**Fig 7A, B, C**). After one month of cryo-preservation *LmCen*^*−/−*^ parasites vials were thawed and used directly in animals without further *in vitro* culture. To evaluate the immune response to this GLP-grade parasite, hamsters were immunized with either GLP-grade or laboratory-grade *LmCen*^*−/−*^ parasites. The expression of pro-inflammatory IFN-γ and TNF-α cytokines was determined after stimulation of splenic cells recovered from hamsters at six weeks post immunization (**Fig. 7D, E**). Both GLP-grade *LmCen*^*−/−*^ and laboratory-grade *LmCen*^*−/−*^ induced comparable levels of pro-inflammatory IFN-γ and TNF-α cytokines that were significantly higher compared to naïve control hamsters (**Fig. 7D, E**). Furthermore, analysis of spleen and liver parasite loads after six- and nine-months post needle challenge with *L. donovani* resulted in equivalent control of parasitemia in animals immunized with either GLP-grade or laboratory-grade parasites that was significantly reduced in comparison to non-immunized control hamsters (**Fig. 7F, G**). Taken together these data validate the immunogenicity of GLP-grade *LmCen*^*−/−*^ parasites that induce a similar protective immunity as laboratory-grade parasites against visceral infection in a pre-clinical animal model.

**Fig. 7.**
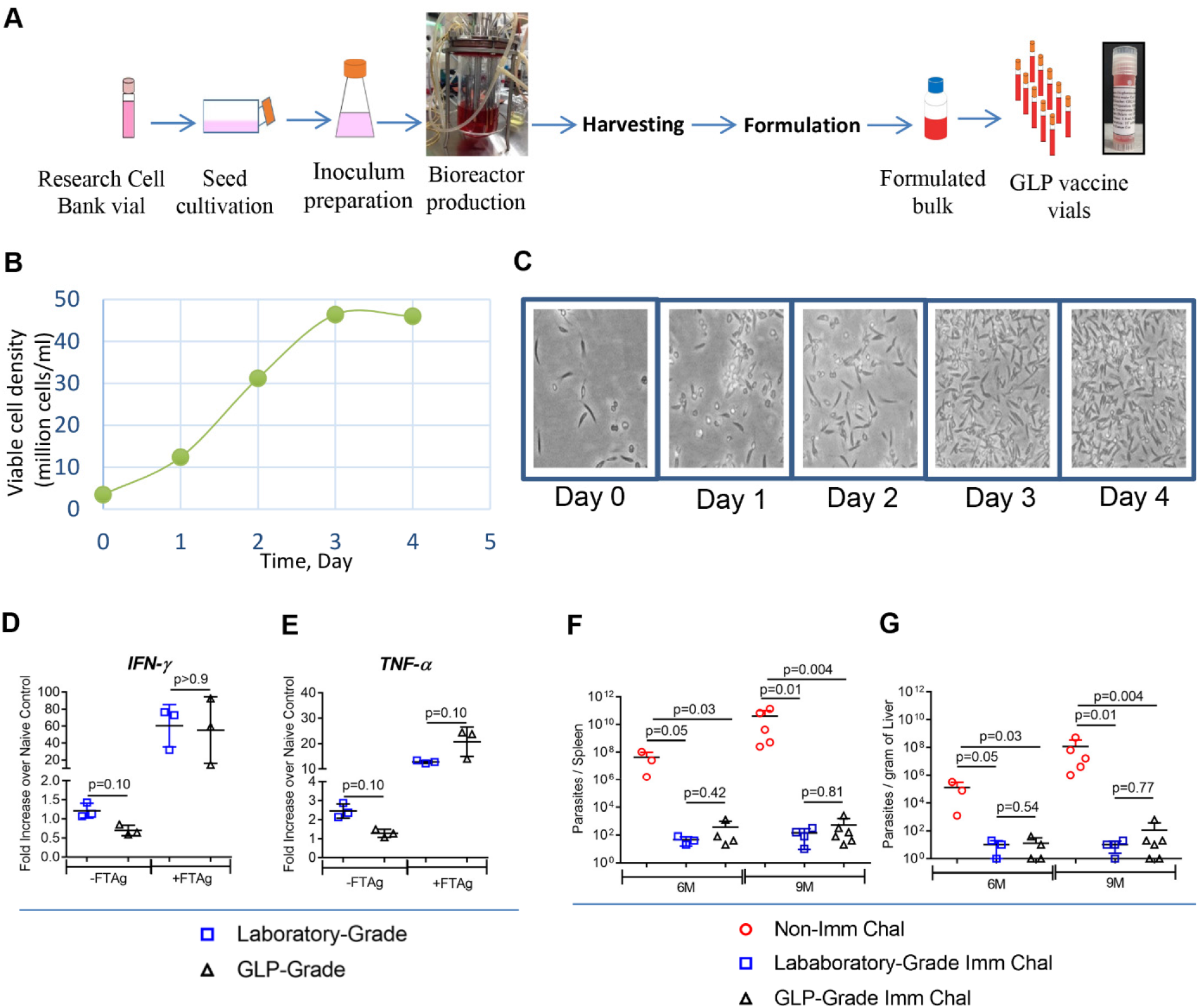
GLP-grade *LmCen*^*−/−*^ parasites induce host protection in hamsters against wild type infection. (**A, B, C**) Production of GLP-grade *LmCen*^*−/−*^ parasites. (A) Seed train of bioreactor production process starting with cell bank vial and culturing into T-flasks. (B) Growth behavior of parasites during bioreactor cultivation, (C) Morphology (magnification 400x) of parasite during four days in bioreactor. (**D** and **E**) Expression of INF-γ (D) and TNF-α (E) in the spleen (with or without 24h of *L. major* freeze thaw antigen re-stimulation; ±FTAg) of hamsters following 6 weeks post immunization with either laboratory-grade (n=3) or GLP-grade (n=3) *LmCen*^*−/−*^ parasite. Results were normalized to γ-Actin expression and shown as the fold change relative to age matched naive hamster. Results (Mean ± SD) are representative of one experiment (p values were determined by Mann Whitney two-tailed test). (**F** and **G**) Parasite loads in the spleen (F) and liver (G) of hamsters either immunized with laboratory-grade (Laboratory-Grade Imm Chal, n=3-4) or GLP-grade (GLP-Grade Imm Chal, n=4-6) *LmCen*^*−/−*^ parasites or age matched non-immunized control (Non-Imm Chal, n=3-5) were determined by limiting dilution following 6 and 9 months post needle challenge with *L. donovani* and expressed as number of parasites per Spleen and per gram of Liver. Results (Mean ± SD) are representative of one experiment (p values were determined by Mann Whitney two-tailed test).

### GLP-grade *LmCen*^*−/−*^ parasites induce pro-inflammatory cytokines in human PBMCs

Next, we evaluated the nature of the immune response induced by GLP-grade *LmCen*^*−/−*^ in PBMCs from healthy individuals from the USA, and from healthy and cured VL subjects from Bihar state, India, non-endemic and endemic for VL, respectively, (**Fig. 8**). The levels of IFN-γ and IL-10, key pro-inflammatory and anti-inflammatory cytokines implicated in resistance or susceptibility to VL, respectively, were measured from culture supernatants of PBMCs 48h after infection with GLP-grade *LmCen*^*−/−*^ parasites. Uninfected PBMCs from the same subjects were used as controls. Induction of IFN-γ and IL-10 following *LmCen*^*−/−*^ infection was significantly higher in healthy-non-endemic (**Fig. 8A)**, healthy-endemic (**Fig. 8B**) and cured VL (**Fig. 8C)** groups compared to the response of their respective control uninfected PBMCs. Further, pairwise analysis revealed elevated expression of IFN-γ in PBMCs from all subjects in the two healthy groups, and the cured VL group (Fig. S5A, B, C). For IL-10, pairwise analysis revealed that all healthy non-endemic and cured VL individuals, and six of seven healthy-endemic individuals showed elevated expression of IL-10. However, it is important to note that levels of IL-10 were ~ two-fold lower compared to IFN-γ with the ratio of IFN-γ to IL-10 being significantly higher in the *LmCen*^*−/−*^ group compared to the non-infected group in all cohorts (**Fig. 8A, B, C**).

**Fig. 8.**
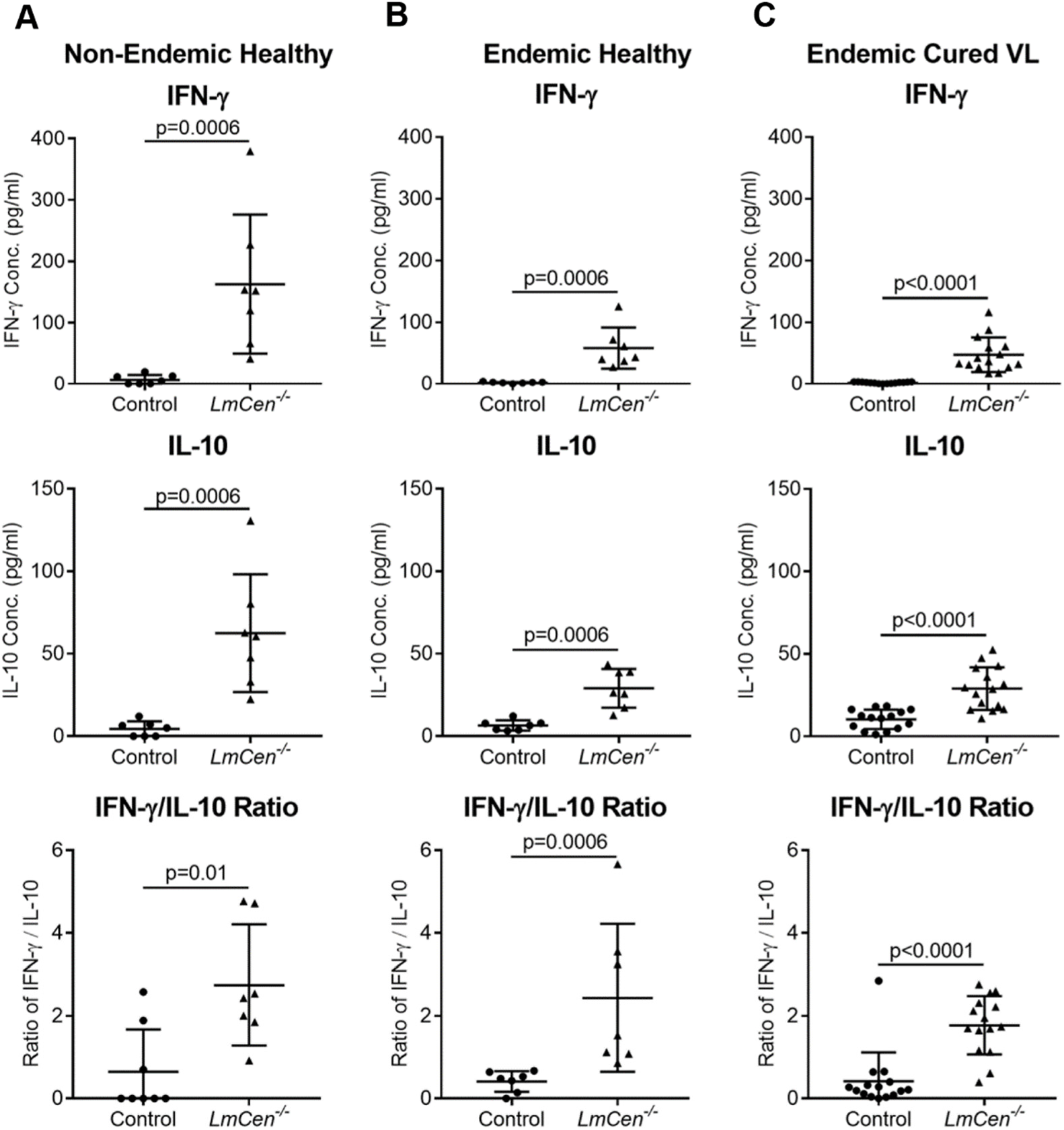
GLP-grade *LmCen*^*−/−*^ parasites induce pro-inflammatory cytokines in the PBMC’s from healthy human subjects living in Nonendemic, VL endemic region and Cured VL patients. (**A, B, C**) Scatter dot plots showing the levels (pg/ml) of pro-inflammatory (IFN-γ) and anti-inflammatory (IL-10) cytokine as well as IFN-γ / IL-10 ratio in culture supernatant of PBMCs from Non-endemic and endemic healthy people (n=7) and cured VL patient (n=15) after 48h of infection with *LmCen*^*−/−*^ parasites by ELISA. Control group was left un-infected. Results (Mean ± SD) are representative of one experiment (p values were determined by Mann-Whitney two tailed t test).

## DISCUSSION

In this study, we demonstrate that vaccination with a dermotropic *LmCen*^*−/−*^ protects against sand fly-initiated fatal visceral leishmaniasis in hamsters caused by the viscerotropic *L. donovani* parasite. We have previously shown that immunization with *LmCen*^*−/−*^, a dermotropic *L. major*, can protect against homologous infection in mouse models (***30***). It has been previously reported that infection with some *Leishmania* species can confer cross-protection against different species of *Leishmania* (***24, 27, 28***).

We showed that this vaccine is safe in tested animals. Intradermal immunization with the live attenuated *LmCen*^*−/−*^ did not develop any signs of disease pathology at the inoculation site compared to wild type *L. major* infection, even under immune-suppressed conditions in the tested animals. However, the persistence of a low number of *LmCen*^*−/−*^ parasites, detected only following immune-suppression, may be important for maintaining long term protection as reported in other studies (***32, 33***).

We also showed that vaccination with *LmCen*^*−/−*^ will not result in sand flies transmitting this parasite as xenodiagnoses experiment demonstrates that *LmCen*^*−/−*^ parasite do not survive inside the sand fly gut, thus non-transmissible from immune person to non-immune person. Furthermore, experimental evidence suggests genetic exchange can happen between different *Leishmania* species in the sand fly vector (***34, 35***), which may lead to the reversion of attenuated parasites to virulent form. This event is unlikely to happen as *LmCen*^*−/−*^ parasites do not survive inside the sand fly gut.

Our result suggests that *LmCen*^*−/−*^ immunization develops a pro-inflammatory environment in both the tissues with an abundant expression of Th1 cytokines. The ear tissue and antigen-stimulated spleen cells from immunized animals significantly up-regulate the expression of pro-inflammatory cytokines IL-1β and IFN-γ, along with TNF-α and IL-12p40. In contrast, IL-10 expression, a major regulatory cytokine plays a crucial role in disease progression either by inhibiting the development of Th1 cells (***36***) or by blocking macrophage activation by IFN-γ (***37***), was significantly down-regulated in the vaccinated animals. Further, the efficacy of a vaccine candidate depends on the ratio of IFN-γ and IL-10 (***38***). A significantly higher ratio of IFN-*y* to IL-10 was observed in the vaccinated animals than in the wild type infected animals predictive of the protective potential of this live attenuated parasite. In addition, a lower expression of anti-inflammatory cytokines IL-4 and IL-21 in the vaccinated group further corelates with induction of host protective immunity and control of future infections (***7, 39***). The inter-relationship between transcription factors T-bet and STAT1, GATA3 and STAT6 and Foxp3, master regulators of Th1, Th2 and T-regulatory cell (Treg) development respectively, also determine the host immune response and out-come of the disease (***40, 41***). Significant up-regulation of T-bet, and STAT1 with concomitant down-regulation of GATA3, STAT6 and Foxp3 in antigen re-stimulated spleen cells, suggests a biased Th1 phenotype in the vaccinated animals. Interleukin-6, a pleiotropic cytokine, showed elevated expression in both the ear, the site of vaccination, and in antigen stimulated spleen cells, correlating with previous study demonstrating that IL-6 controls *Leishmania* disease progression by reducing the accumulation of regulatory T cells (***42***). Further, the increased expression of IL-1β, IL-6, IFN-γ, STAT1, CCL5 and CXCL9 genes associated with M1 polarization of macrophages, along with the down-regulation of Arg-1, CCL17 and CCL22 genes, a characteristic of M2 polarization of the macrophages (***43***), indicates that *LmCen*^*−/−*^ immunization biases towards the M1 phenotype in vaccinated animals, and suggests that macrophages play an important role in the *LmCen*^*−/−*^ mediated immune response to control parasitemia.

We observed that *LmCen*^*−/−*^ immunization induced significant host protection as early as six weeks of post-challenge with wild type *L. donovani* parasites. Further, parasite replication was significantly controlled afterwards compared to non-immunized challenged animals using needle challenge. Because the protective efficacy of a vaccine against needle challenge is not always equally effective compared to vector challenge (***44, 45***), we validated the vaccine efficacy of *LmCen*^*−/−*^ in a natural sand fly infection model. *LmCen*^*−/−*^ induces significant protection against *L. donovani* infected sand fly challenge, as evidenced by the robust control of parasitemia both in the spleen and liver up to 12-month post challenge. Furthermore, All the immunized-challenged hamsters remained healthy compared to non-immunized challenged hamsters in both infection models. This protection was associated to a Th1 response observed in the ear and by antigen re-stimulated spleen cells in the *LmCen*^*−/−*^ immunized hamsters following *L. donovani* challenge with needle injection, or systemically from antigen re-stimulated spleen cells after sand fly challenge. Importantly, *LmCen*^*−/−*^ immunized animals exhibited a significantly higher ratio of IFN-*y* to IL-10 as reported in cured VL patients as a potential regulatory mechanism to control visceral infections (***38, 46***).

To explore whether *LmCen*^*−/−*^ parasites as a vaccine candidate we grew *LmCen*^*−/−*^ parasites under GLP conditions, later on this process can be produced in cGMP compliant facility for clinical trials. We demonstrated that GLP-grade *LmCen*^*−/−*^ parasites are equally immunogenic and protective as laboratory grown parasites against wild type *L. donovani* infection in a preclinical hamster model. Moreover, GLP-grade *LmCen*^*−/−*^ parasites induced IFN-γ in PBMCs of healthy human subjects living in non-endemic or VL endemic regions, indicative of priming towards a Th1-biased immune response. Corroborating our data, a recent study demonstrated that an anti-leishmanial DNA vaccine induced an IFN-γ-dominated immune response in healthy human volunteers (***11***). Further, dominance of IFN-γ over IL-10 in PBMCs from healthy or VL-cured individual upon exposure to GLP-grade *LmCen*^*−/−*^ parasites confirmed their induction of a predominantly pro-inflammatory Th1-type immune response, a pre-requisite for an effective *Leishmania* vaccine. Together, these data demonstrate the pre-clinical safety and protective efficacy of a GLP-grade *LmCen*^*−/−*^ parasite vaccine that is ready to be tested in a first-in-humans clinical trial.

There are two perceived limitations to this study. One, although hamsters have been used as the gold standard animal model for VL, because of its similarity to humans in the outcome of mortality and morbidity, analysis of various parameters are constrained by the lack of availability of a broad range of immunological reagents for hamsters. Secondly, the immunological response of human samples from endemic region, presented in this study, is limited to PBMCs stimulated with GLP-grade material, and needs to be validated in clinical trials using GMP-grade material. We are currently in the process of manufacturing cGMP grade *LmCen*^*−/−*^ parasites and are planning for a Phase 1 and Phase 2 clinical study of the *LmCen*^*−/−*^ vaccine in endemic regions of VL to assess its safety and explore immunological correlates of protection.

To our knowledge, this is the first report of a marker free dermotropic genetically modified *Leishmania* vaccine that is protective against a viscerotropic *Leishmania* validated *in vivo* against a natural vector challenge in a preclinical animal model, and that induces IFN-γ *ex vivo* in PBMCs of human subjects. Taken together, this study indicates that *LmCen*^*−/−*^ is safe, efficacious against VL and could be explored as vaccine candidate by clinical trial in both endemic as well as nonendemic of VL.

## MATERIALS AND METHODS

### Study design

In this study we wanted to determine the vaccine efficacy of *centrin* gene deleted *LmCen*^*−/−*^ parasites against experimental VL in hamsters. All animal experimental procedures used in this study were reviewed and approved by the Animal Care and Use Committee of the Center for Biologics Evaluation and Research, U.S. Food and Drug Administration and the National Institute of Allergy and Infectious Diseases (NIAID) (http://grants.nih.gov/grants/olaw/references/phspolicylabanimals.pdf). The number of animals used in this study vary from 6-15 hamsters per group.

### Animals and Parasites

Six to eight-week-old female outbred Syrian golden hamsters (*Mesocricetus auratus*) were obtained from the Harlan Laboratories (Indianapolis, IN) and housed either at the Food and Drug Administration (FDA) animal facility, silver spring (MD) or National Institute of Allergy and Infectious Diseases (NIAID) Twin-brook animal facility, Rockville (MD) under pathogen-free conditions. The wild type *L. donovani* (*LdWT*) (MHOM/SD/62/1S) parasites, wild type *L. major* Friedlin (FV9) (*LmWT*) and *centrin* gene deleted *LmCen*^*−/−*^ (Friedlin strain) promastigotes were cultured as previously described (***30, 47***).

### Hamster immunization and determination of parasite load

Six to eight-week-old female hamsters were immunized with 10^6^ total stationary-phase either laboratory-grade or GLP-grade *LmCen*^*−/−*^ parasites by intradermal injection in the ear in 10μl PBS using a 29-gauge needle (BD Ultra-Fine). Control group of animals were infected with 10^6^ total stationary-phase *L. major* wildtype (*LmWT*) promastigotes. Lesion size was monitored by measuring the diameter of the ear lesion using a direct reading Vernier caliper. Parasite burden in the ear, draining lymph node (dLN), spleen and liver was estimated by limiting dilution analysis as described in previous studies (***47***).

### Xenodiagnosis

Hamsters infected with *LmWT* and *LmCen*^*−/−*^ parasites were xenodiagnosed at 2- and 8-weeks post infection. Briefly, five to seven-day old unfed female *Lutzomyia longipalpis* sand flies allowed to feed on the inoculated ear of anesthetized hamsters (the site of inoculation of *LmWT* & *LmCen*^*−/−*^ parasites) for 1 hour in the dark. Blood-fed sand flies were separated and maintained in chambers under controlled conditions for eight days. Midgut dissections were carried out and examined by microscopy for the presence of live *Leishmania* parasites.

### Immunosuppression by dexamethasone treatment

To determine the safety of *LmCen*^*−/−*^ parasites in immune-suppressive condition, 6- to 8-weeks old hamsters were divided into three groups. Group-1 (n=6) were infected with 10^6^ stationery phase *LmWT* parasites and Group-2 (n=12) and Group-3 (n=12) animals were immunized with 10^6^ stationery phase *LmCen*^*−/−*^ parasites in a 10μl volume of PBS through intradermal routes (into the ear dermis). After 10-weeks of post infection, only Group-3 animals were treated with 2 mg/kg Dexamethasone sodium phosphate (Sigma Aldrich) in PBS by subcutaneous injection three times, alternate days for one week. Four weeks after this treatment (total 15-weeks post infection), all the animals in three different groups were sacrificed and evaluated for parasite load by serial dilution as described (***47***). Development of pathology & lesion size in the ear was assessed at 15-weeks post inoculation by measuring the diameter of the lesion.

Characterization of *centrin* gene deleted parasites isolated from *LmCen*^*−/−*^ immunized and DXM treated group was done by Polymerase chain reaction. Total Genomic DNA was isolated from the parasites recovered from immune-suppressed hamsters as well as *LmWT* and *LmCen*^*−/−*^ parasites according to the manufacturer information (DNeasy Blood & Tissue Kit, Qiagen). PCR was performed with *L. major centrin* gene specific primer (For-5’-ATGGCTGCGCTGACGGATGAACAGATTCGC-3’; Rev-5’-CTTTCCACGCATCTGCAGCATCACGC-3’) which target the amplification of the 450-bp. A reaction mixture was prepared containing 10× Buffer (Invitrogen), 0.2mmol/l each deoxyribonucleotide (Invitrogen), 1μmol/l each primer, 1.25-units of Taq-polymerase (Invitrogen) and 200ng of DNA samples in a final volume of 50μl. The PCR conditions were as follows: denaturation at 94°C for 3min, followed by 35 cycles of 94°C for 20s, 58°C for 20s and 68°C for 35s with a final extension of 68°C for 5min. The amplification reactions were analyzed by 1% agarose gel-electrophoresis, followed by ethidium bromide staining and visualization under UV light. DNA from the reference *plasmid* (PCR 2.1 TOPO) containing *centrin* gene was used as a positive control.

### Cytokine determination by real-time PCR

Cytokine expression in hamster’s tissue (ear and spleen) were determined by real-time PCR at indicated time points. Briefly, total RNA was extracted using PureLink RNA Mini kit (Ambion). Aliquots (400 ng) of total RNA were reverse transcribed into cDNA by using random hexamers from a high-capacity cDNA reverse transcription kit (Applied Biosystems). Cytokine gene expression levels were determined by either TaqMan probe (TaqMan, Universal PCR Master Mix, Applied Biosystem) or SYBR green (Applied Biosystems) PCR using a CFX96 Touch Real-Time System (BioRad, Hercules, CA). The sequences of the primers (forward and reverse) and probes (5′ 6-FAM and 3′ TAMRA Quencher) used to detect the gene expression are shown in Table-1 (Table S1) (***43, 48***). The data were analyzed with CFX Manager Software. The expression levels of genes of interest were determined by the 2^−ΔΔCt^ method; samples were normalized to either γ-actin or 18S rRNA expression and determined relative to expression values from naive hamsters.

### Immunizations and needle challenge studies

Six to eight weeks old female hamsters were immunized with 10^6^ stationery-phase either laboratory-grade or GLP-grade *LmCen*^*−/−*^ parasites in a volume of 10μl PBS through intradermal routes. After 7 weeks of immunization, the animals were needle challenged (ID route) with virulent 5×10^5^ meta-cyclic *Ld1S* promastigotes into the contralateral ear. As a control group, age-matched non-immunized hamsters were similarly challenged with virulent 5×10^5^ meta-cyclic *Ld1S* promastigotes. After various periods of post challenge (1.5-, 3-, 6-, 9- & 12-month post challenge), animals were sacrificed & organs (spleen and liver) were removed aseptically, weighed and parasite load was measured by limiting dilutions as previously described (***47***). Hamsters were monitored daily during study, and their body weights were recorded weekly. As an additional confirmation for the presence of parasites in tissue, multiple impression smears from spleen were prepared and examined for the parasites.

### Sand Fly Challenge studies

Female 2- to 4-day-old *Lutzomyia longipalpis* sand flies were infected by artificial feeding on defibrinated rabbit blood (Spring Valley Laboratories, Sykesville, MD) containing 5 × 10^6^/ml *L. donovani* amastigotes supplemented with 30μl penicillin/streptomycin (10,000 units penicillin/10 mg streptomycin) per ml of blood for 3 h in the dark as described Aslan H et al. (***49***). Parasite loads and percentage of metacyclic per midgut were determined using hemocytometer counts. Sand flies on day 13-after infection were used for subsequent transmission to hamsters. Thirty flies with mature infections were applied to collateral ear of each *LmCen*^*−/−*^ immunized and age matched non-immunized hamsters through a meshed surface of vials held in place by custom-made clamps. During exposure to sand flies, hamsters were anesthetized intraperitoneally with ketamine (100 mg/kg) and xylazine (10 mg/kg). The flies allowed to feed for 2h in the dark at 26°C and 75% room humidity. As a qualitative measure of transmission, the number of blood-fed flies was determined. Each hamster received an average of 15-infected bites per transmission. Hamsters were monitored daily during infection, and their body weights were recorded weekly. After various periods post challenge (3.5-, 9- and 12-month), animals were sacrificed & organs (spleen and liver) were removed aseptically, weighed and parasite load was determined by limiting dilutions method as previously described (***47***). Furthermore, multiple impression smears of spleen tissue were prepared and examined for the presence of parasites in the tissue.

### Manufacturing of GLP-grade *LmCen*^*−/−*^ vaccine

In one-liter bioreactor, modified M199 medium supplemented with 10% FBS was used to produce GLP-grade *LmCen*^*−/−*^ parasites. To inoculate 1L bioreactor, seed cultivation was started from cell bank. One cell bank vial was thawed and transferred entirely into T-25 cm^2^ flask having 5 ml of medium. After 2-3 days, culture was expanded into T-75 cm^2^ flask having 15ml of medium. Further, after 2-3 days, culture was transferred into shake flask for inoculum development. After 2-3 days of shake flask cultivation, 1L bioreactor was inoculated containing 500ml medium. Bioreactor run was done for four days where culture reached its maximum cell density of 40-50 × 10^6^ cells/ml. During bioreactor cultivation, parasites were found healthy and highly motile. At day 4, about 80-94hr when culture was in early stationary phase, parasites were harvested by centrifugation. Harvested parasites were washed with PBS and formulated to make vaccine. GLP vaccine vials were filled with formulated bulk where one vaccine vial contained 1.8 ml of 10 × 10^6^ cells/ml of live attenuated parasites.

### Human Blood donors and ethical consideration

We randomly recruited a total of 7- endemic healthy volunteers (male/female, 4:3; age range, 21-60 years) and 15-treated VL patients (male/female, 9:6; age range, 22-65 years) for this study. Endemic healthy individuals were recruited after confirmation of no previous report of VL/PKDL infection, negative rK39 ICT status and no current clinical symptoms of VL (fever, anemia, and splenomegaly). All the cured VL patients had recovered from VL infection by treatment with anti-leishmanial drug, viz. Ambisome or Miltefosine and were discharged from the hospital about 6-months before their recruitment in this study. They had no clinical symptoms of VL and were positive for rK39 ICT. Exclusion criteria included HIV or any other co-infection like tuberculosis, pneumonia etc.; other diseases including malaria, hepatitis B, and hepatitis C; use of any immunosuppressive drugs, with history of relapse of VL, and pregnant or lactating females.

The protocols of human study on Indian blood donors have been reviewed and approved by institutional ethical committee of partner institution ICMR-Rajendra Memorial Research Institute of Medical Sciences, India (RMRI/EC/26/2019). Written informed consent was obtained from each participant of the study. PBMCs of seven different healthy donors from non-endemic region were collected from ATCC, USA (PCS-800-011™).

### PBMC isolation and infection

10 ml of peripheral venous blood from each cured VL patients and healthy control volunteers (endemic) was collected in LPS-free conditions into Li-heparin vacutainers (BD Biosciences, SanJose, CA, USA). The PBMC was isolated within 10 min after blood collection by density gradient centrifugation over Ficoll-Paque Plus (GE Healthcare, Piscataway, NJ, USA) according to the manufacturer’s instructions. Briefly, 3ml of Ficoll-Paque PLUS was added to a 15 ml centrifuge tube and 4 ml of blood sample was carefully over-layered on that. The tube was then centrifuged at 900xg for 30 min at 20°C. After centrifugation, the PBMC layer (comprised of lymphocytes/monocytes), generated in the middle of plasma and Ficoll-Paque PLUS, was carefully aspirated out and placed in a fresh tube. The isolated cells were then washed twice with sterile PBS (Invitrogen) at 300 g (5 min at 20°C). The viability of the PBMC was >98%, as checked by trypan blue dye exclusion method.

PBMCs were re-suspended in complete RPMI-1640 medium (Invitrogen), supplemented with 100 U/ml penicillin (Invitrogen), 100 mg/ml streptomycin (Invitrogen), and 20 mM Hepes, pH 7.4 (Sigma-Aldrich), and plated at a density of 2×10^6^ cells/ml in tissue culture dishes (Nalge Nunc Int., Rochester, NY, USA). These plates were incubated for 72 hours for adherence at 37°C in 5% CO_2_. After this time, the autologous non-adherent cells were slowly aspirated out from the first plate and transferred to a second fresh plate and kept at 37°C in 5% CO_2_.

Fresh complete RPMI-1640 medium was immediately added to the adherent cells in the first plate and kept at 37°C in 5% CO_2_. The adherent cells were then either kept uninfected or were infected with *LmCen*^*−/−*^. After 6 hours post-infection, the unbound parasites were removed by washing twice with PBS. After washing, non-adherent autologous cells in the second plate were added back to the first plate and co-cultured with the uninfected/infected adherent cells for 48-hour time points at 37°C in 5% CO_2_. The supernatant was collected after the incubation, centrifuged at 400g and the cell-free fraction was stored at −80° C for cytokine ELISA.

### Histological staining

For histology, liver from hamsters were fixed in fixative solutions (10% buffered formalin phosphate solution). Sectioning and hematoxylin and eosin (H&E) staining of all samples was done by Histoserv (Gaithersburg, MD). Stained sections were analyzed under the microscope (KEYENCE).

### Statistical analysis

Statistical analysis of differences between groups was determined by unpaired two-tailed Mann-Whitney or Wilcoxon pairs two tailed t test, using Graph Pad Prism 7.0 software. The heat map of the gene clusters was assembled in GraphPad Prism 7.0 software, using the log10 values of relative fold expression of cytokines, chemokines and transcription factors over naive.

## Author contributions

SK (Subir Karmakar), NI, FO, JO, WWZ, SK (Swarnendu Kaviraj), KPS, AM, SD, KP, MS, SS, RS, SG, PB, GV and RD conducted experiments, analyzed data and helped to write the manuscript. SK (Subir Karmakar), RD, AS, SH, PD, SK (Shaden Kamhawi), SS, JGV and HLN designed experiments, analyzed data and wrote the manuscript.

## Data availability statement

The data that support the findings of this study are available from the corresponding author upon reasonable request.

## Materials & Correspondence

Correspondence and requests for material should be addressed to H.N., or R. D.

## Supplementary Figures

**Fig. S1.**
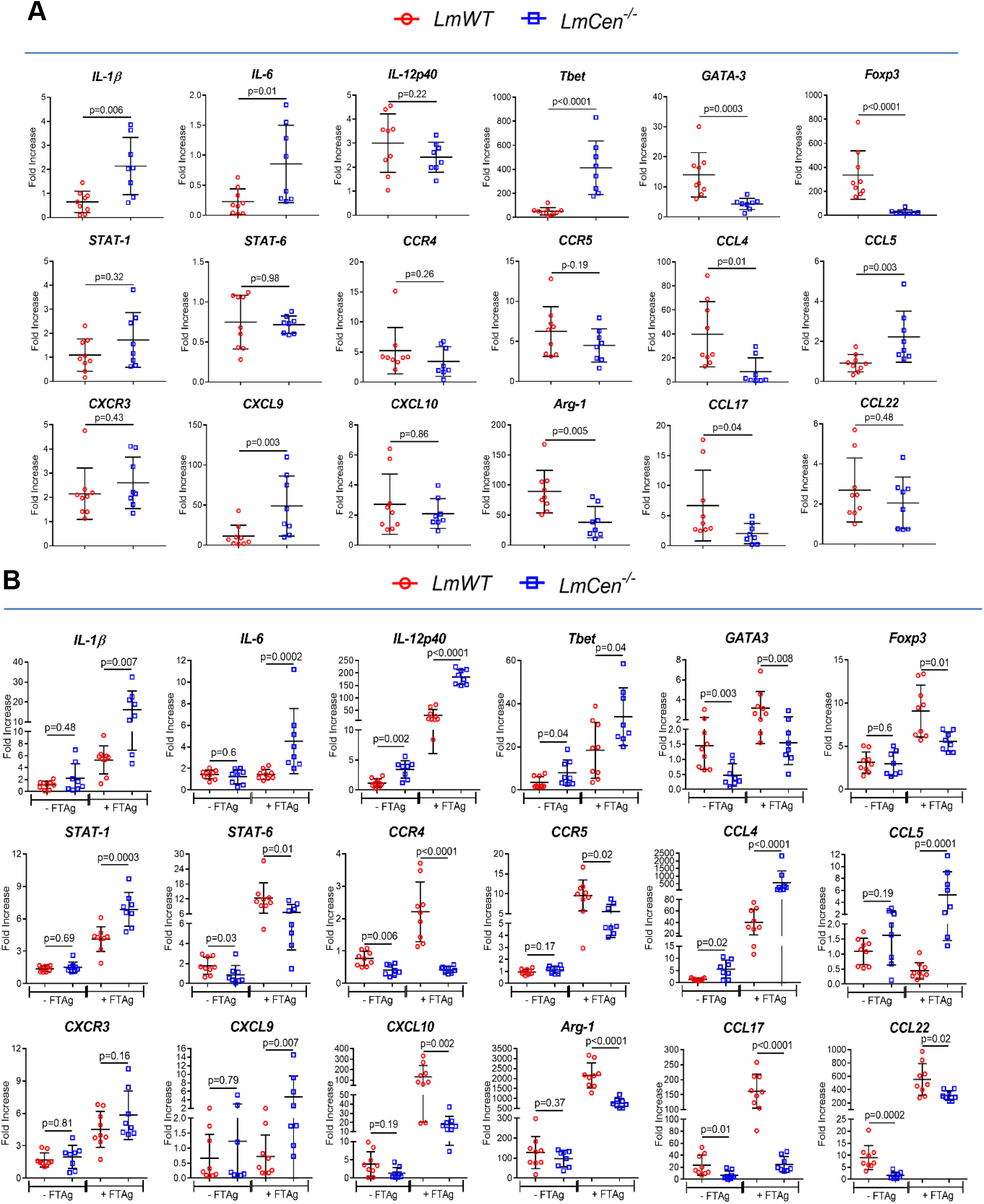
*LmCen*^*−/−*^ immunization induces pro-inflammatory immune response. (**A** and **B**) The expression of transcripts (IL-1β, IL-6, IL-12p40, T-bet, GATA3, Foxp3, STAT-1, STAT-6, CCR4, CCR5, CCL4, CCL5, CXCR3, CXCL9, CXCL10, Arg-1, CCL17 and CCL22) was confirmed by RT-PCR in the inoculation site (ear) (A) (without antigen re-stimulation) and spleen (B) (with or with-out 24h of *L. major* freeze thaw antigen re-stimulation; ±FTAg) of hamsters immunized with *LmCen*^*−/−*^ parasites (n=8) and compared with *LmWT* (n=9) infected group following 7 weeks of post inoculation. The data were normalized to either γ-Actin or 18S expression. Results (mean ± SD) are representative of cumulative effect of two independent experiments. Statistical analysis was performed by non-parametric Mann-Whitney two-tailed test.

**Fig. S2.**
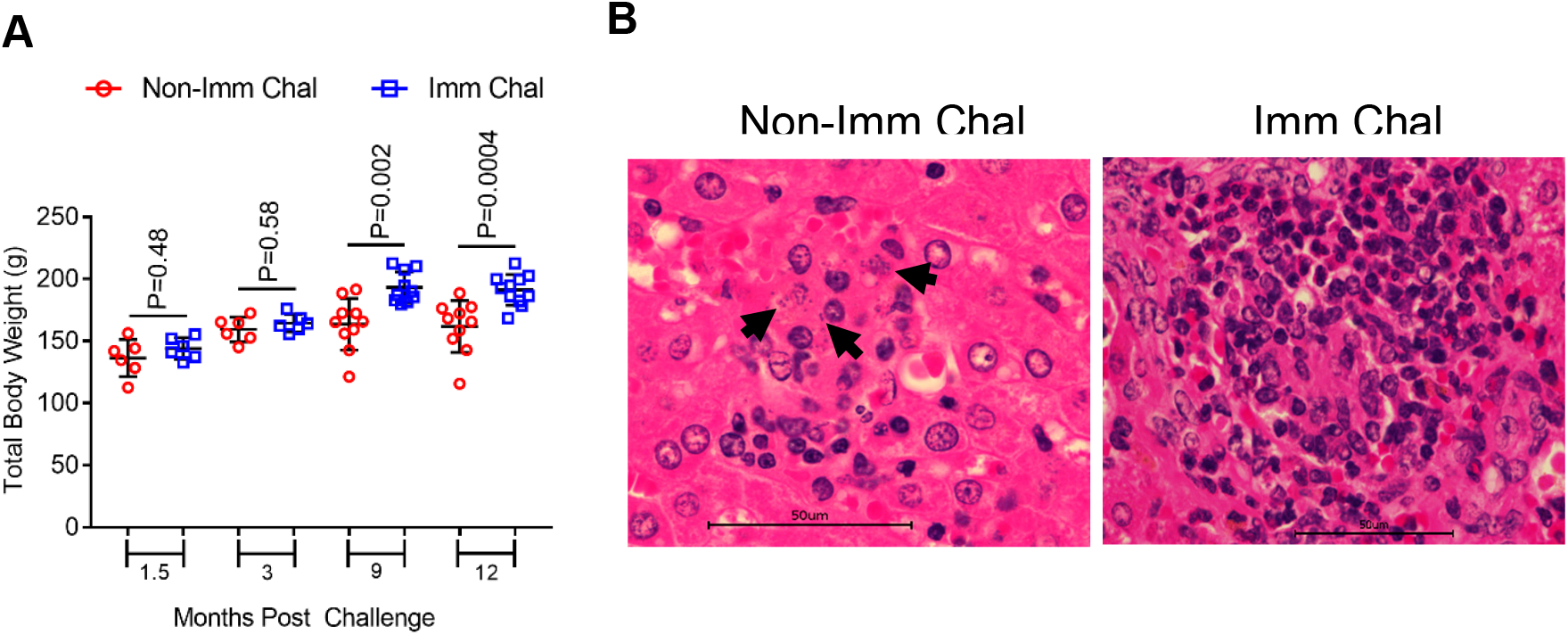
Clinical outcomes of *LmCen*^*−/−*^ immunized hamsters following needle challenge with virulent *L. donovani*. (**A**) Body weight of *LmCen*^*−/−*^ immunized (Imm Chal) (n=6 for 1.5 and 3 months and n=10 for 9 and n=11 for 12 months) and age matched non-immunized (Non-Imm Chal) (n=6 for 1.5 and 3 months and n=10 for 9 and 12 months) hamsters after various periods of post needle challenge (1.5, 3, 9 & 12 month) with *L. donovani.* Results (mean ± SD) are representative of cumulative effect of two independent experiments. Statistical analysis was performed by Mann Whitney two-tailed test. (**B**) Photomicrograph of hepatic histology from one of the representative non-immunized (Non-Imm Chal) and immunized ((Imm Chal) hamster at 9-months after *L. donovani* needle challenge. Liver section from immunized animal shows well-developed mature granuloma free of parasites with surrounding mononuclear infiltrated cells. In non-immunized animal, granuloma is less organized with heavily parasitized kupffer cells (black arrow) and few infiltrating lymphocytes.

**Fig. S3.**
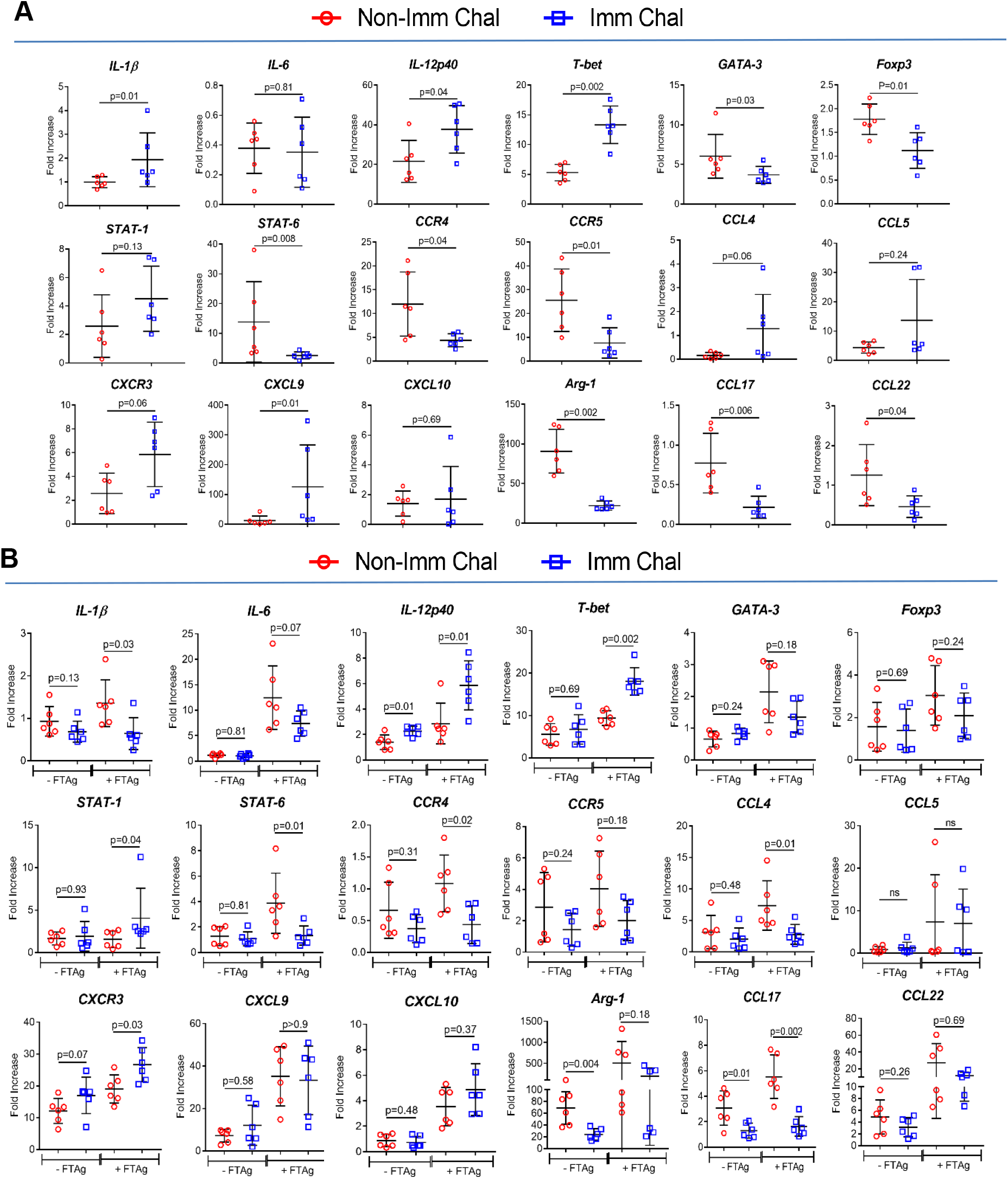
*LmCen*^*−/−*^ immunized hamster induces pro-inflammatory/Th1 type of immune response upon challenge with wild type *L. donovani*. (**A** and **B**) The expression of transcripts (IL-1β, IL-6, IL-12p40, T-bet, GATA3, Foxp3, STAT-1, STAT-6, CCR4, CCR5, CCL4, CCL5, CXCR3, CXCL9, CXCL10, Arg-1, CCL17 and CCL22) was confirmed by RT-PCR in the challenged ear (A) (without antigen re-stimulation) and spleen (B) (with or with-out 24h of *L. donovani* freeze thaw antigen re-stimulation; ±FTAg) of *LmCen*^*−/−*^-immunized (Imm Chal, n=6) and age matched non-immunized (Non-Imm Chal, n=6) hamsters at 1.5 months of post *L. donovani* needle challenge. The data were normalized to either γ-Actin or 18S expression. Results (mean ± SD) are representative of cumulative effect of two independent experiments. Statistical analysis was performed by non-parametric Mann-Whitney two-tailed test.

**Fig. S4.**
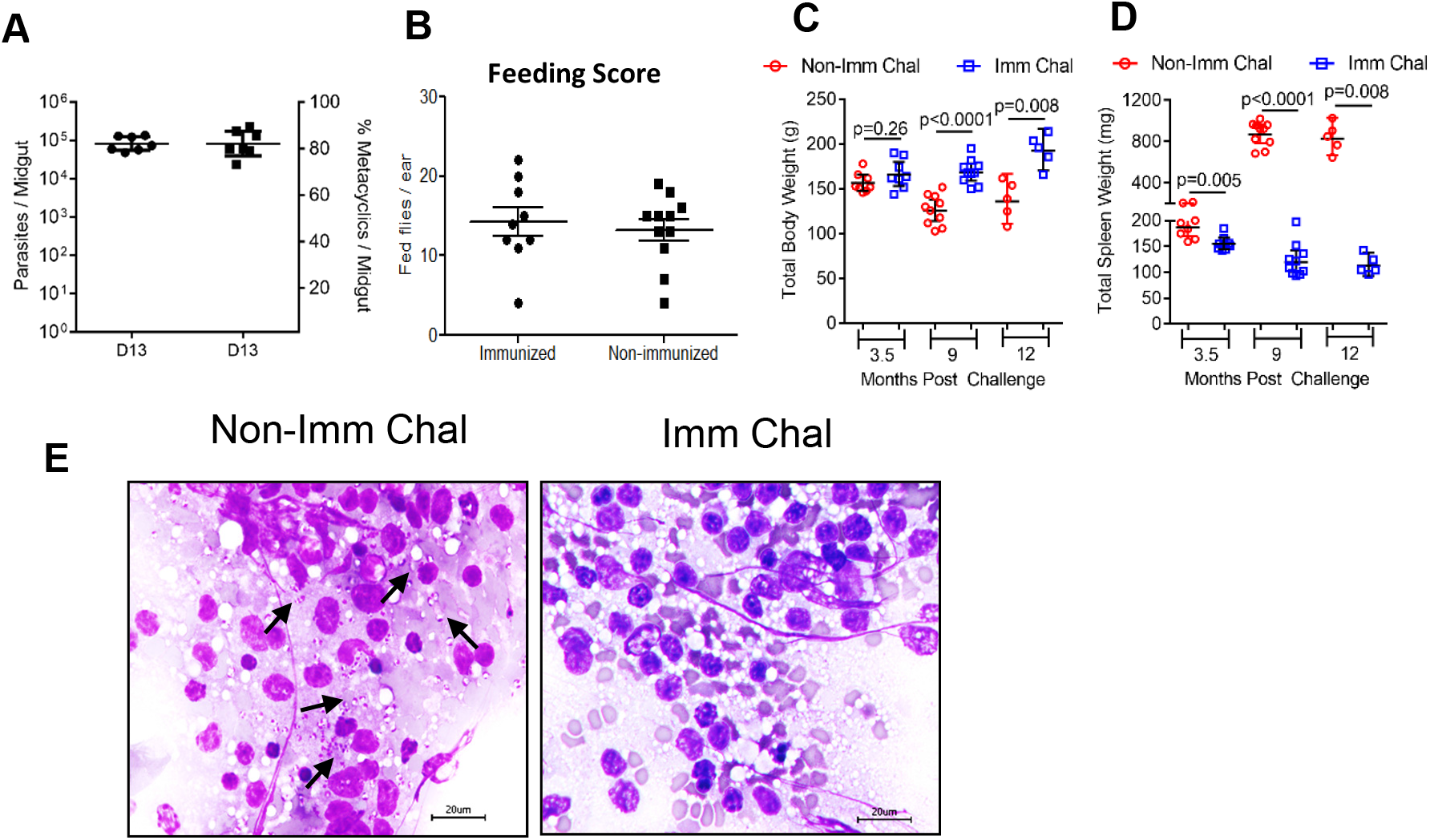
Clinicopathology of *LmCen*^*−/−*^ immunized/non-immunized hamsters after different time of post *L. donovani* infected sand fly challenge. (**A**) Assesment of the parasite load and percent metacyclics per sand fly after 13 days of post sand fly infections. For challenge studies, 13 days of post infected sand flies were used. (**B**) Post transmission feeding score from 30 total sand flies exposed to right ears of each hamster. Data from one of two independent experiments are shown. (**C** and **D**) Seven weeks *LmCen*^*−/−*^ immunized (Imm Chal)/age matched non-immunized hamsters (Non-Imm chal) were challenged into the contralateral by the bite of *L. donovani*-infected sand flies (*Leutzomia longipalpis*). After various periods (3.5, 9 & 12 month) of post challenge, animals were sacrificed and following parameters were observed (C) Body weight (g) and (D) spleen weight (mg). Results are representative of cumulative effect of two independent experiments for 3.5- and 9-months post-challenge and one experiment for 12 months post-challenge. Bars represent the geometric means with 95% Cl of total 5-10 hamsters (n=8, n=10 and n=5 for 3.5M, 9M and 12M post challenge respectively of non-immunized and immunized challenged groups) in each group are shown. Statistical analysis was performed by non-parametric Mann-Whitney two-tailed test. (**E**) Tissue impression smear stained with H&E from spleen, showing intense parasitism of a representative non-immunized (Non-Imm chal) hamster compared to *LmCen*^*−/−*^ immunized (Imm Chal) hamster at 12-months of post *L. donovani* infected sand fly challenge. Black arrow showing the amastigotes in the impressions. Bar = 20 μm.

**Fig. S5.**
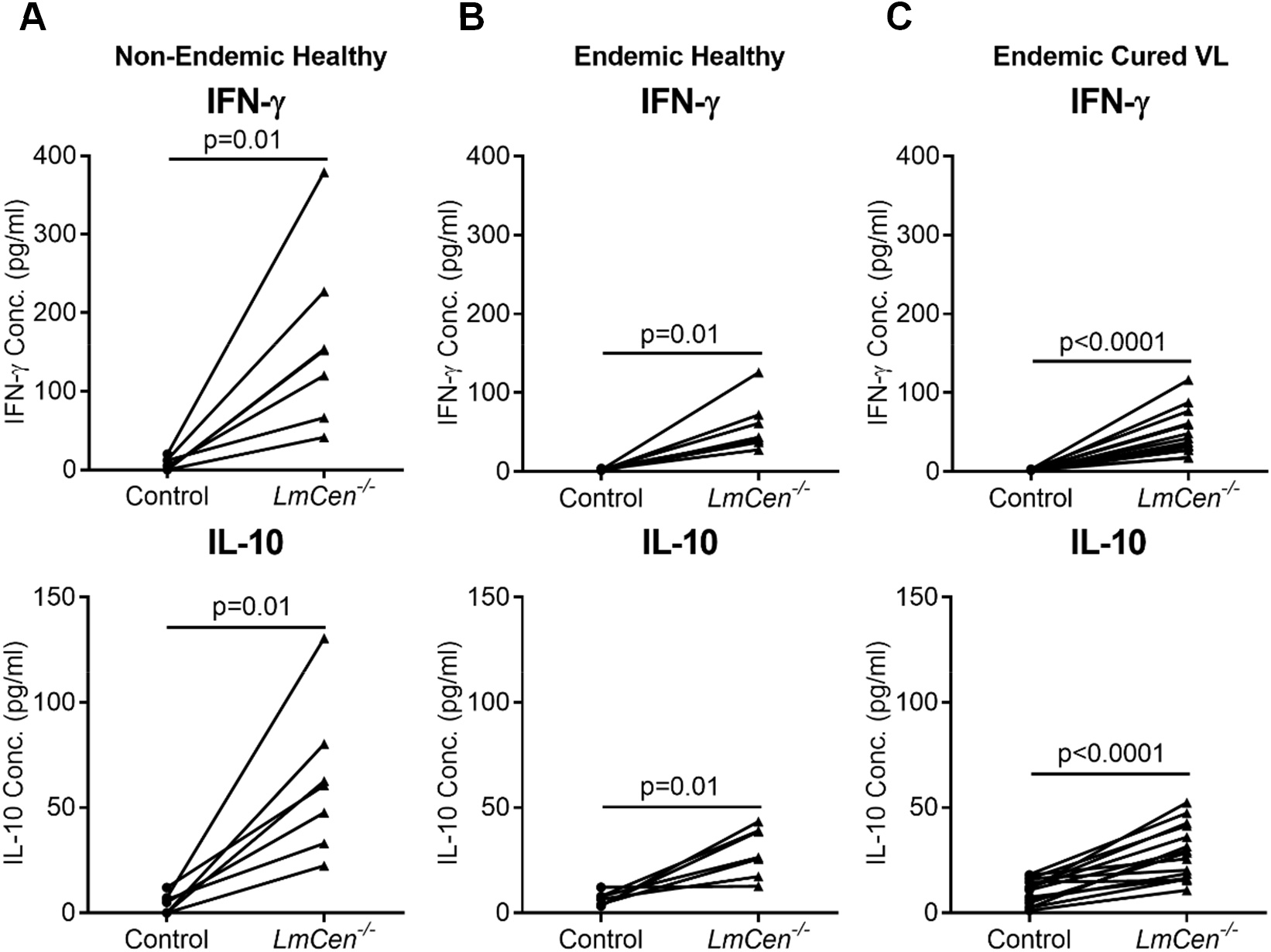
GLP-grade *LmCen*^*−/−*^ parasites induce pro-inflammatory cytokines in the Human PBMC’s. (**A, B, C**) Ex-vivo pairwise analysis of pro-inflammatory (IFN-γ) and anti-inflammatory (IL-10) cytokine levels from culture supernatant of PBMC’s from healthy human subjects living in Non-endemic (A), VL endemic region (B) as well as from cured VL patients (C) of endemic region after 48h of infection with *LmCen*^*−/−*^ parasites. Control group was left un-infected. A statistically analysis of paired samples (n=7 for healthy individuals from non-endemic and endemic region and n=15 for cured VL patients) for each cytokine were represented. Significance was determined by Wilcoxon pairs two tailed t test.

**Table S1:**
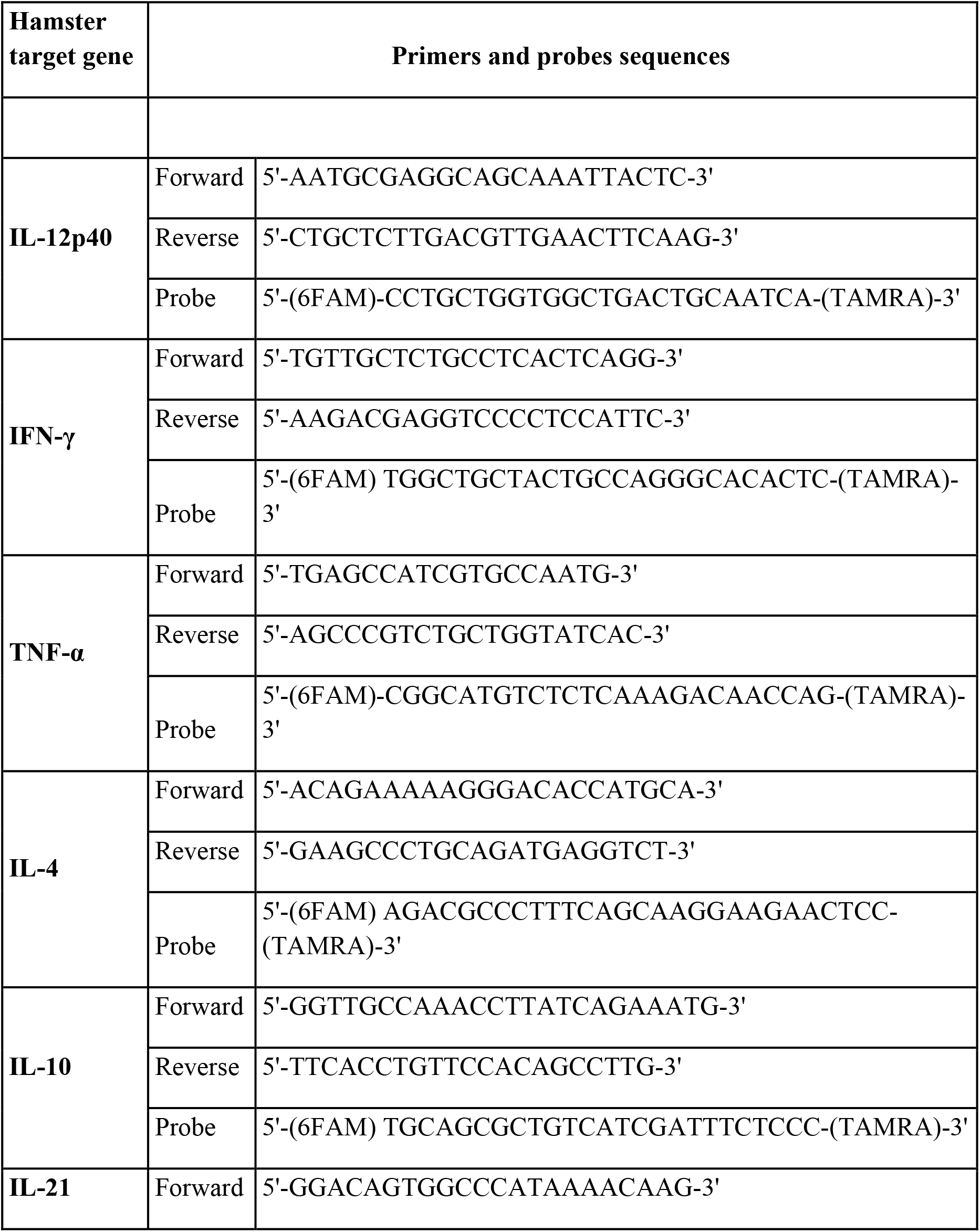
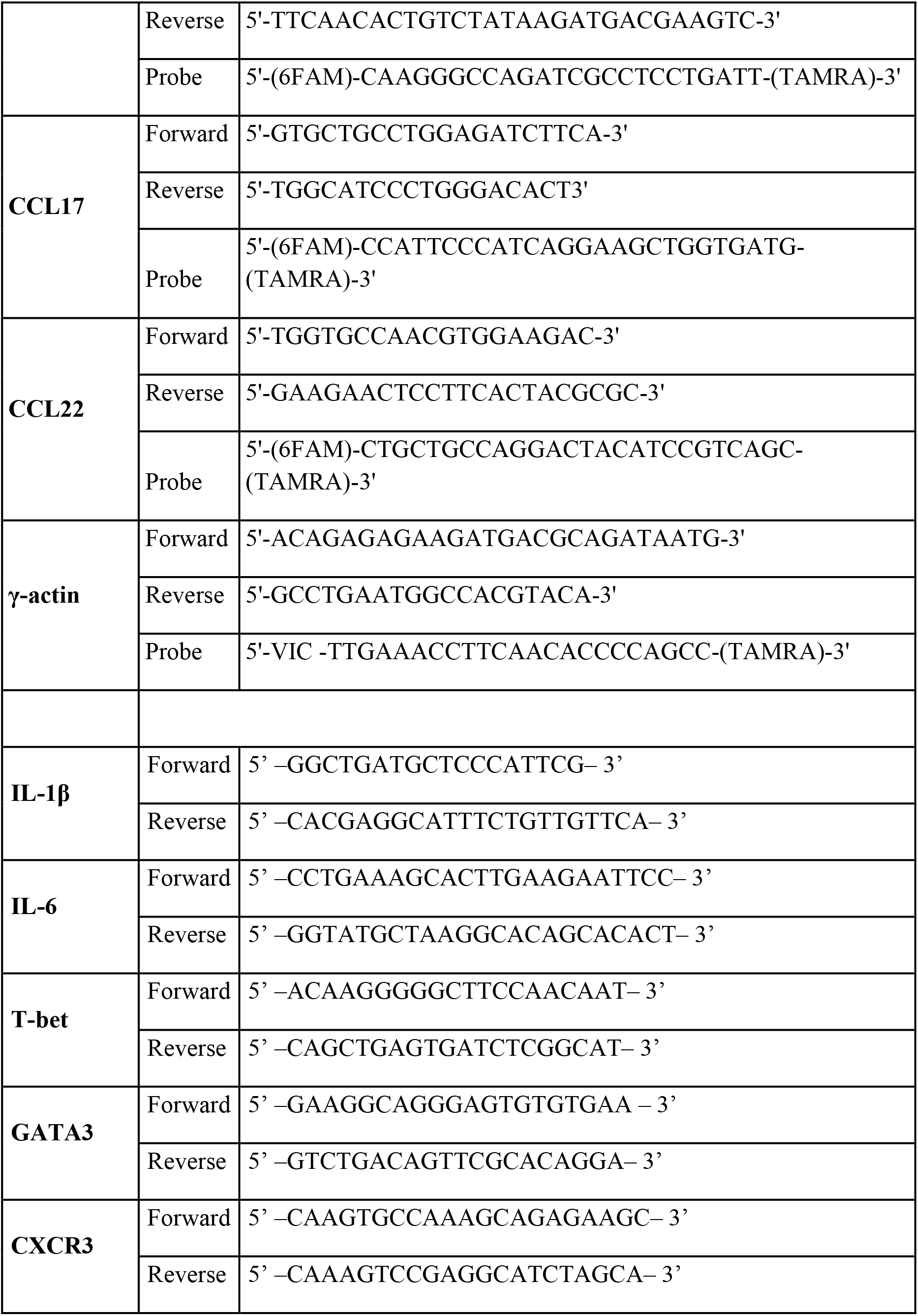
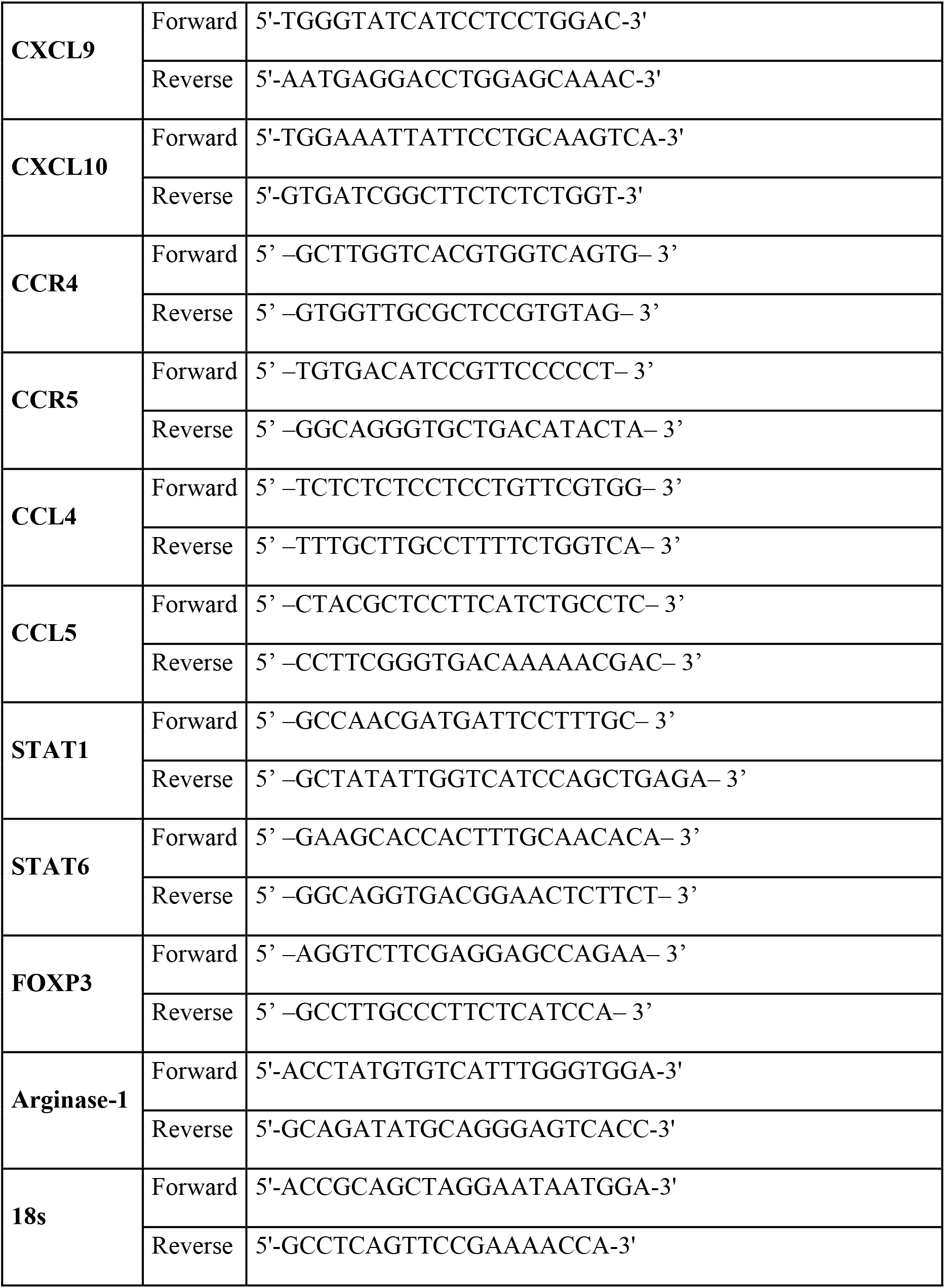
Primers and probe sequences.

